# Molecular Structure, DNA Binding, and Photophysical Properties of SYTOX Orange and SYTOX Green

**DOI:** 10.64898/2026.07.08.737150

**Authors:** Koen R. Storm, Stefanie D. Pritzl, Yi-Yun Lin, Christian Wiebeler, Alptuğ Ulugöl, Martin Lehmann, Dave J. van den Heuvel, Gerhard A. Blab, Gerd Gemmecker, Jan Lipfert

## Abstract

Fluorescent dyes are critical to visualizing nucleic acids in many applications. SYTOX Orange and SYTOX Green are cyanine dyes, used in dead cell staining and increasingly in single-molecule assays to probe DNA supercoiling and processing. However, their structures and effects on DNA mechanics are not or only partially known.

We determine the structure of SYTOX Orange to be (E)-2-((2-(4 ((diethyl(methyl)ammonio)methyl)phenyl)-6-methoxy-1-methylquinolin-4(1H)-ylidene)methyl)-4-methyloxazolo[4,5-b]pyridin-4-ium, identical to SYBR Gold except for an aza-benzoxazol core that is fundamentally different from other dyes in the SYTOX and SYBR families. We report SYTOX Green to be (Z)-2-(bis(3-(trimethylammonio)propyl)amino)-4-((3-methylbenzo[d]thiazol-2(3H)-ylidene)methyl)-1-phenylquinolin-1-ium, similar to PicoGreen. Using magnetic tweezers, we characterize the effect of SYTOX Orange and SYTOX Green on DNA mechanics. They lengthen and unwind DNA consistent with intercalation and the DNA unwinding angles per dye are 21.1(1)° and 20.5(1)° for SYTOX Orange and Green, respectively. Both dyes leave the DNA bending persistence length and plectoneme size almost unaltered (<10% change up to 1 µM), which is advantageous in assays probing DNA supercoiling. Their photophysical properties reveal close agreement between single-molecule manipulation and optical absorbance and fluorescence spectroscopy. Our comprehensive set of complementary measurements relates mechanical and optical properties to the molecular structures and provides recommendations for their use in applications.

## INTRODUCTION

The interaction of DNA with ligands is fundamental for many cellular processes as well as biotechnological applications. In particular, fluorescent dyes are routinely used to label DNA for visualization and quantification in a wide variety of assays ranging from imaging of cells, the analysis and quantification of gel bands or PCR products, to single-molecule assays for visualization and manipulation of DNA ^1, 2, 3^.

Phenanthridinium dyes have been used extensively as DNA intercalators and DNA and RNA-fluorescent markers (ethidium bromide) and as probes for cell viability (propidium iodide). However, the more modern cyanine dyes (including the SYBR and SYTOX families) offer both structural and optical tunability, allowing a range of substituent replacements and heterocycle selection to tune their emission range ^2, 4, 5^.

SYTOX dyes are popular modern dead-cell stains and commonly employed in microbiology using flow cytometry, fluorescence microscopy, and spectroscopy ^6^. First introduced with SYTOX Green (SxG), the modern membrane-impermeable fluorescent cyanine dye shows improved performance over the more classical viability dye propidium iodide ^4, 6, 7, 8, 9, 10^. The series of SYTOX dyes has since expanded to offer a large spectral range, facilitating the application of multicolor assays ^5^. Their fast kinetics require only short incubation times, high quantum yield enhancement (>100-fold increase in fluorescent quantum yield upon intercalation into DNA ^11, 12^) results in low background fluorescence, and their high photostability enables long measurements ^13, 14^. More recently, SYTOX dyes have been used in single-molecule assays, in applications ranging from polarization microscopy of DNA ^15^ to real-time monitoring of DNA-protein interactions and processing ^16, 17, 18, 19, 20^. Specifically, SYTOX Orange (SxO) has been used in single-molecule assays to directly visualize DNA supercoiling and plectonemes ^21, 22^ and to observe the action of DNA processing enzymes ^13, 16, 17, 18, 20, 23, 24^.

Despite its widespread use, many important aspects of SxO- and SxG-DNA binding are currently unknown or poorly understood. In particular, it is unclear how SYTOX binding alters the bending stiffness or plectoneme size of DNA, which is important to know whenever SYTOX-stained DNA is used to study DNA dynamics and processing. Further, the angle by which SxO or SxG intercalation unwinds DNA is unknown, which is an important parameter to model supercoil and topology dependent binding and genomic processing ^25, 26, 27, 28^.

Here we use nuclear magnetic resonance (NMR) and mass spectrometry (MS) to resolve the molecular structures of SxO and SxG. We use single-molecule magnetic tweezers (MT) measurements to directly monitor the mechanical response of DNA upon SxG and SxO intercalation. In MT, DNA molecules are tethered between a surface and small (∼1 µm diameter) magnetic beads. External magnets allow to control both the applied stretching forces ^29, 30, 31, 32, 33^ and the DNA rotation, i.e. its linking number ^29, 34^. Using MT, it is possible to probe both the stretching and twist response of DNA as a function of intercalator concentration ^35, 36, 37, 38, 39^. We complement the single-molecule MT measurements with optical spectroscopic measurements of SxO and SxG in the absence and presence of DNA. Furthermore, we characterize the optical properties of these dyes via quantum chemical calculations.

Our NMR and MS investigation shows that SxO is a cyanine dye with an unusual aza-benzoxazol containing core and an overall structure close to SYBR Gold and SxG a thiazole orange derivative similar to PicoGreen. Our MT data find that binding of SxO and SxG to DNA lengthens and unwinds the DNA helix, as expected for intercalative binding. We determine the unwinding angle per intercalated dye to be *Δθ* = 21.1° ± 0.1° for SxO and *Δθ* = 20.5° ± 0.1° for SxG, comparable to values found for similar intercalators. Importantly, we find that SxO and SxG intercalation has only minor effects on the DNA bending persistence length and size of the plectonemes for the concentration range typically employed (≤ 1 µM), which makes SxO and SxG attractive for assays that aim to visualize and probe DNA supercoils and their processing by enzymes through staining. Our optical spectroscopic investigation of SxO and SxG in the absence and presence of DNA finds binding parameters fully consistent with the single-molecule MT measurements. We observe a bathochromic (i.e. to longer wavelengths) shift of the absorbance as well as a dramatic (≥ 1000-fold) increase in fluorescence emission upon intercalation. Our data provide a baseline for modeling DNA in the presence of SxO and SxG and suggest recommendations for its use as a stain for DNA visualization. In addition, our results for the SYTOX dyes enable a comparison to other popular DNA stains such as PicoGreen and SYBR Gold and aid in the selection of dyes for specific applications, as well as offer critical information on fluorophore design.

## RESULTS

We carry out a comprehensive characterization of SxO and SxG and their interactions with DNA using several complementary experimental techniques. We determine the molecular structures of both dyes with a combination of NMR and MS. We then use single-molecule MT measurements to monitor intercalation and to determine the effects of SxO and SxG on the mechanical properties and structure of DNA. Finally, we present absorption and fluorescence spectroscopy data to quantitate the photophysical properties of both the free and DNA-bound dyes and rationalize observed trends using quantum chemical calculations.

### Structure determination by NMR spectroscopy

From NMR analysis the molecular structure of SxO was determined as (E)-2-((2-(4-((diethyl(methyl)ammonio)methyl)phenyl)-6-methoxy-1-methylquinolin-4(1H)-ylidene)methyl)-4-methyloxazolo[4,5-b]pyridin-4-ium (Figure 1a, Supplementary Table S1, Supplementary Figures S1 and S2, and Materials and Methods). At room temperature several ^1^H and ^13^C resonances in the quinoline and aza-benzoxazole parts showed severe line broadening, probably caused by hindered rotation at the methine bridge (Supplementary Figure S1a). NMR spectra required for assignment (especially the ^1^H,^13^C-HMBC long-range correlations) had to be recorded at higher and lower temperatures where the broadened resonances became narrower. The assignment of all ^1^H, ^13^C and ^15^N signals is given in Supplementary Table S1. The SxO core structure consists of a 4-methyl-4-azabenzoxazole and quinoline heterocycle connected by a monomethine group; it is identical to the structure of SYBR Gold except for the modified benzoxazole moiety.

**Figure 1.**
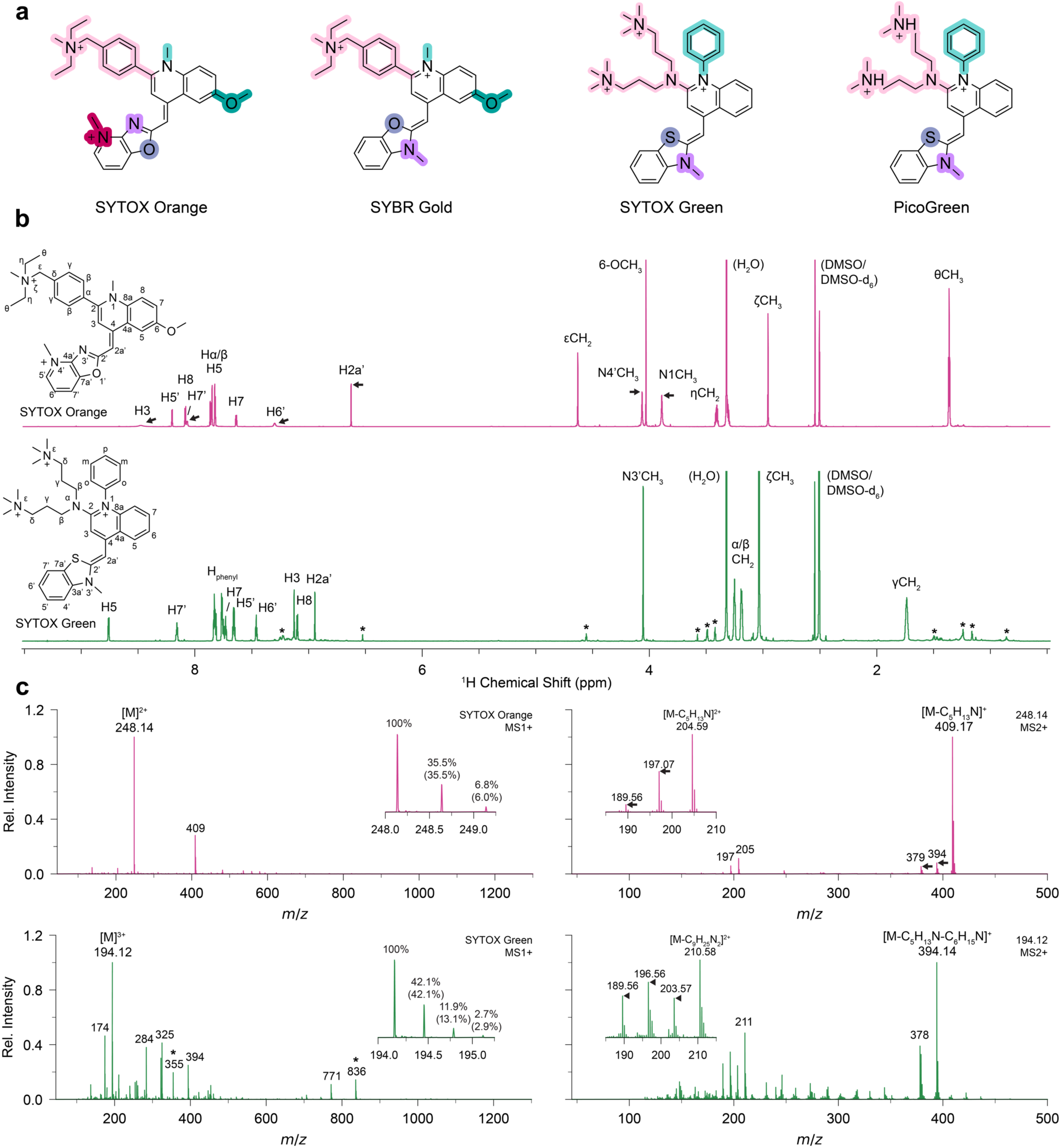
Structure determination of SYTOX Orange and SYTOX Green. a) The structures of SYTOX Orange and SYTOX Green determined in this work by NMR and mass spectrometry. For comparison, the closely related structures of SYBR Gold ^39^ and PicoGreen ^91^ are shown. b) ^1^H NMR spectra of SYTOX Orange (top) and SYTOX Green (bottom) in DMSO-d_6_ (1200 MHz, 298 K) with assignment (see also Supplementary Table 1). Arrows indicate exchange-broadened signals, asterisks indicate signals from impurities. c) ESI MS/MS of SYTOX Orange (top) and SYTOX Green (bottom) in DMSO/MeOH. MS1 are dominated by the intact molecules (*m*/*z* 248.14 for SxO and *m*/*z* 194.12 for SxG). Inset shows the isotope abundance with calculated values in brackets. MS2 spectra of the intact molecules are dominated by fragmentation of the aliphatic side group. Arrows indicate subsequent CH_3_ loss, arrowheads indicate CH_2_ loss, asterisks indicate intact complexes with iodide.

Similarly, we determined the structure of SxG to be (Z)-2-(bis(3-(trimethylammonio)propyl)amino)-4-((3-methylbenzo[d]thiazol-2(3H)-ylidene)methyl)-1-phenylquinolin-1-ium (Figure 1a, Supplementary Table 1, and Supplementary Figure S1 and S2). The full ^1^H, ^13^C and ^15^N assignments are given in Supplementary Table S1. The SxG core structure consists of a benzothiazole and quinoline heterocycle connected by a monomethine group. It is identical to the structure of PicoGreen except for two additional N-methyl groups at the end of the aliphatic quinolone sidechain, forming two positively charged quaternary amines.

### Mass spectrometry of SxO and SxG

To confirm the molecular mass of SxO and SxG and to provide information on substituents and charge states, we performed mass spectrometric analyses. The MS acquired in positive mode reveal a large contribution of the intact dye molecules, at *m*/*z* 248.14 with *z* = 2 for SxO and *m*/*z* 194.12 with *z* = 3 for SxG. MS acquired in negative mode reveal the presence of iodine with *m/z* 126.90 as the anion for both dyes (Figure 1c and Supplementary Figure S3). The exact masses of 496.28 u for SxO and of 582.36 u for SxG from the MS analyses are in line with the approximate values of ∼500 u and ∼600 u, respectively, reported by the manufacturer ^5^ and in excellent agreement with the masses of the predicted structures of 496.28 u and 582.36 u. The mass spectra of SxO reveal a pattern of fragments that is very similar to SYBR Gold ^39^, with peaks shifted by exactly Δ(*m/z*) *= –*1, which confirms that both dyes have essentially identical substituents to the core fluorophore scaffold, and suggests a substitution in the core ring system. Similarly, the isotope distribution of the compounds (Figure 1, insets) shows close agreement with the calculated values (percentages in brackets). The relatively large M+2 peak for SYTOX Green can be attributed to sulfur in its benzothiazole group. The MS/MS of the intact molecules are dominated by fragmentation of the aliphatic side groups. Fragmentation of SxO appears to predominately be loss of its (diethyl(methyl)ammonio)methyl side chain and subsequent loss of CH_3_, while for SxG cleavage in its (trimethylammonio)propyl side chains is apparent together with various losses of CH_2_.

### Magnetic tweezers measurements

To determine the effects of SxO on the structure and mechanical properties of DNA, in particular to monitor intercalation induced lengthening and unwinding of the DNA helix, we performed single-molecule MT measurements (Figure 2; Materials and Methods). In a first set of experiments, we measured the DNA extension at different levels of applied force for torsionally relaxed DNA. In a second set of measurements, we probed the response of DNA to systematic over- and underwinding at constant forces. For all MT measurements, we kept the DMSO concentration in the buffer to ≤ 2% to avoid effects due to the presence of DMSO ^40^ (Supplementary Figures S4 and S5) and flushed ≥ 25 cell volumes when changing dye concentration to ensure equilibration (Supplementary Figures S6 and S7).

**Figure 2.**
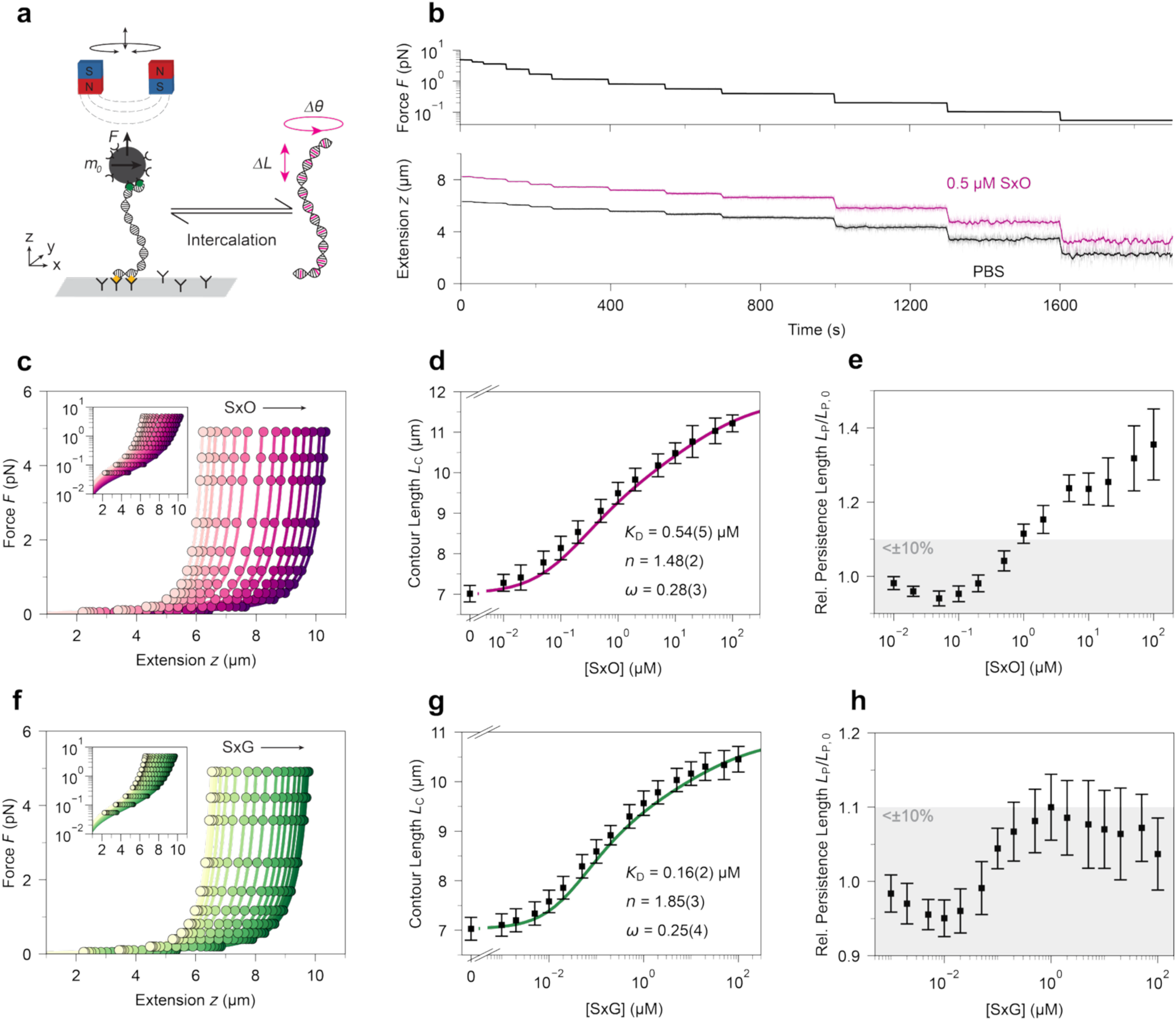
Magnetic tweezers force-extension measurements probe changes in DNA contour length and bending persistence length with SxO and SxG concentration. a) Schematic of the MT measurements. DNA molecules are tethered between a glass surface and magnetic beads in a flow cell. Permanent magnets above the flow cell enable the application of stretching forces and torques, respectively. Intercalation into DNA causes lengthening and changes in its twist. b) Extension time traces for 20.6 kbp DNA in PBS (black) and in 0.5 μM SxO (red). The DNA extension (bottom) is measured while the stretching force is systematically varied (top). c) Force-extension curves for DNA in the presence of increasing SxO concentrations in PBS. The concentrations are (from light pink to purple) 0, 0.01, 0.02, 0.05, 0.1, 0.2, 0.5, 1, 2, 5, 10, 20, 50, 100 μM SxO. Solid lines are fits of the worm-like chain (WLC) model. d) DNA contour length *L*_C_ as a function of SxO concentration obtained from WLC fits as shown in panel c. The solid line is a fit of the cooperative McGhee–von Hippel model to the force extension and rotation-extension data (Figure 2). Values shown are the dissociation constant, binding site size and cooperativity factor from the fit. e) DNA bending persistence length *L*_P_ as a function of SxO concentration as obtained from WLC fits as shown in panel c. f-h) Same as panels c-e but for increasing SxG concentration (from light yellow to green, 0.001-100 μM). Data in panels d, e, g, and h are the mean ± s.d. from at least 8 independent molecules. The grey shading in panels e and f indicate changes by less than 10% relative to bare DNA.

### Magnetic tweezers force-extension measurements monitor how SxO and SxG intercalation lengthens DNA

To study the effect of SxO and SxG on DNA length and bending stiffness, we performed force-extension measurements on torsionally unconstrained (i.e. nicked) DNA in MT (Figure 2a,b). DNA force-extension measurements over a broad range of dye concentrations are well described by the inextensible WLC model ^35, 41, 42, 43^ (Figure 2c,f). First, we measured the response to force of DNA in PBS only (Figure 1c, lightest color). The contour length *L*_C_ and persistence length *L*_P_ determined from the WLC model are *L*_C_ = 7.0 ± 0.2 μm and *L*_P_ = 43 ± 5 nm (mean ± s.d. from 16 independent molecules) (Figure 2d,g), in excellent agreement with the expected crystallographic length of 7.0 μm and with previous measurements under the same conditions ^30, 35, 40^. We then recorded force-extension curves in presence of increasing SxO and SxG concentrations (Figure 1b,c,f). The contour lengths obtained from fitting the WLC model show a systematic increase with increasing SxO and SxG concentrations (Figure 2d,g).

### The binding of SxO and SxG to DNA is well described by the anti-cooperative McGhee-Von Hippel model

The contour lengths determined from MT force-extension measurements systematically increase with dye concentration, by ≈50% compared to bare DNA, characteristic of intercalation. However, we do not observe a clear saturation level; instead, the contour length still increases with dye concentrations, albeit slowly, at the highest tested concentrations, consistent with previous data for SxO (Supplementary Figure S8) and suggestive of anti-cooperative binding^44^. Fits of the non-cooperative McGhee-Von Hippel model ^35, 45^ capture the binding approximately, but fail to describe the behavior at the highest concentrations (Supplementary Figure S9). Consequently, we fit the (anti-)cooperative McGhee-Von Hippel model ^45^ to the change in contour length (and, simultaneously, to the twist of DNA, see below) to obtain the dissociation constant *K*_D_, the binding site size *n*, the cooperativity parameter *ω*, and unwinding angle Δ*θ* (Figure 2d,g; Equations 1,3, and 4). We assume a length increase of Δ*z* = 0.34 nm per intercalated molecule, a value typical for intercalators ^13, 35, 46, 47^. The model can account for binding over the entire concentration range probed for both SxO and SxG (Figure 2d,g, parameters given in the figure legend) and the fitted values of the cooperativity parameter *ω* = 0.28 ± 0.03 for SxO and 0.25 ± 0.04 for SxG are indicative of moderate anti-cooperativity. Anti-cooperative binding of DNA intercalators has been previously described for ethidium bromide ^44^. An elevated off-rate at high dye coverage has been reported for SYBR Gold, YOYO and POPO, likely due to electrostatic repulsion ^13^, suggesting that dye-dye interactions along the DNA result in a reduced effective binding affinity. The fitted dissociation constants of *K*_D_ ≈ 0.5 µM for SxO and ≈ 0.15 µM for SxG are in good agreement with previous reports ^13^ of 0.42 and 0.13 µM, respectively, in particular when taking into account the salt dependence ^13^. Similarly, the fitted values for the binding sites size *n* ≈ 2, indicating intercalation approximately every other base pair at saturation, are in good agreement with previous reports for SxO and SxG ^13^ and with the general observation that intercalator exhibit nearest neighbor exclusion.

### SxO and SxG intercalation have only minor effects on DNA bending stiffness

In contrast to the contour length, which systematically lengthens upon intercalation, the effect on the bending persistence length *L*_P_ varies significantly across different intercalators. For example, intercalation of ethidium bromide systematically decreases the bending persistence length of DNA by ∼1.5-fold ^35, 44^, while binding of YOYO tends to increase it ^48, 49, 50^. We find that addition of low concentrations of SxO or SxG initially leads to a small decrease in *L*_P_ until around 0.1 μM SxO and 0.01 µM SxG, respectively (Figure 2e,h). At higher dye concentrations, *L*_P_ starts to increase slightly with increasing concentrations. An initial decrease followed by an increase of *L*_P_ with increasing dye concentration was previously observed for the closely related dye SYBR Gold ^39^. Despite the observed small variations of *L*_P_ with SxO or SxG concentration, overall *L*_P_ remains fairly constant for the SYTOX dyes. For SxG, the bending persistence changes by less than 10% over the entire concentration range probed (0.001 – 100 µM; Figure 2h), similar to what was previously observed for PicoGreen ^37^. While for SxO concentrations above 1 μM a systematic increase in *L*_P_ is observed (Figure 2e), for both SYTOX dyes *L*_P_ remains essentially within error, changing by ≤ 10% in the concentration range typically used for single-molecule visualization experiments of (<1 µM). A nearly constant bending stiffness upon addition of the intercalator is desirable for assays that use fluorescent intercalators to visualize DNA looping, compaction, and processing, to minimize the perturbation of DNA properties relative to unstained DNA, making the SYTOX dyes an attractive choice for this type of assay.

### Rotation-extension measurements reveal the effect of intercalation on torsionally constrained DNA

In addition to applying forces, MT can control the DNA linking number *Lk* by rotating the magnets. We define as zero turns the point at which DNA in the absence of the intercalator is torsionally relaxed, corresponding to the intrinsic helicity *Lk*_0_ of DNA. Upon over- and underwinding the DNA (corresponding to positive and negative applied turns, respectively) the characteristic supercoiling response of DNA is observed (Figure 3). For experiments at low applied stretching forces (< 1 pN), the effect of the applied turns on the DNA extension is approximately symmetric for positive and negative turns and results in a buckling transition in both directions. Post-buckling, positive or negative plectonemic supercoils are formed in the DNA tether resulting in a linear decrease of the extension with the number of applied turns.

**Figure 3.**
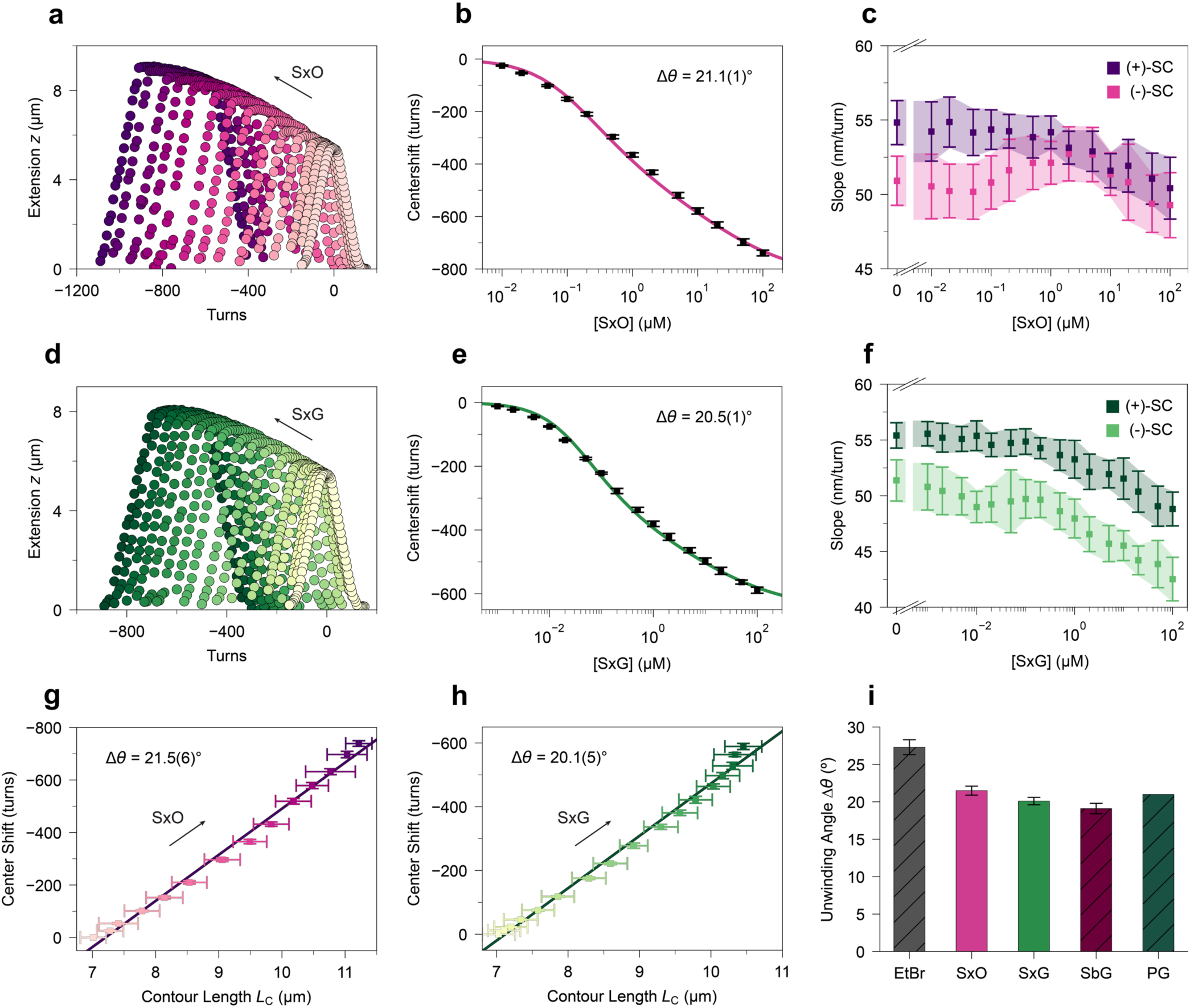
Magnetic tweezers rotation-extension measurements quantitate DNA unwinding by SxO and SxG and their effect on plectoneme size. a) Rotation-extension measurements of a 20.6 kbp DNA in the presence of increasing concentration of SxO at a constant stretching force of *F =* 0.5 pN. The SxO concentrations are (from light pink to purple) 0, 0.01, 0.02, 0.05, 0.1, 0.2, 0.5, 1, 2, 5, 10, 20, 50, 100 μM. b) Quantification of the shift in the center position of the rotation extension curves in the presence of increasing concentration of SxO. The center positions were determined by extrapolation of the slopes in the positive and negative plectonemic regime. The solid line is a fit of the McGhee–von Hippel model to the rotation extension and the force extension data, *Δθ* is the fitted DNA unwinding angle per intercalation event. c) Slopes of the rotation-extension data in the plectonemic regime at *F =* 0.5 pN for positive and negative supercoiling. The slopes provide a measure for the size of the plectonemes and exhibit only a small dependence on SxO intercalation. d-f) Same as panels a-c but for increasing SxG concentration (from light yellow to green, 0.001-100 μM). g) and h) Quantification of the shift in the center position of the rotation-extension curves plotted against the change in contour length from the force extension curves (Figure 2d-g) reveals a linear relation for g) SxO and h) SxG (same concentrations and color code as in panels a-d). From the slope we determine the unwinding angle per SxO to be 21.5° ± 0.6° and per SxG to be 20.1° ± 0.5°. i) Comparison of DNA unwinding angles of SxO and SxG determined in this work with structurally similar intercalators SYBR Gold (SbG) ^39^ and PicoGreen (PG; error bar not shown since it is ±14°) ^37^ and ethidium bromide (EtBr) ^35^. Data in panels b, c, e, and f are the mean ± s.d. from at least 10 independent molecules.

Overall, SxO and SxG have very similar effects on DNA rotation curves and several changes in the rotation extension responses are apparent upon binding (Figure 3a,d). First, the maximum extension increases with increasing intercalator concentration, mostly due to the intercalation induced lengthening discussed in the previous section. Second, a systematic shift to negative turns with increasing concentration occurs, which we quantify by extrapolation of the slopes in the plectonemic regime and determining the intersection points (Supplementary Figure S10). Third, we observe a broadening of the rotation-extension curves with increasing concentration. Fourth, the curves exhibit a slope in the pre-buckling regime, i.e. in the top part of the curves, which becomes more pronounced with increasing concentration. We discuss these changes systematically in the next subsections.

### SxO and SxG intercalation unwinds DNA

The systematic shift of the rotation-extension curves to negative turns with increasing SxO or SxG concentration (Figure 3a,d) is directly related to the unwinding of the DNA helix upon intercalation (Materials and Methods). We employ two different approaches to quantify the unwinding angle per intercalated SxO or SxG molecule from the measured shifts of the extension-rotation curves to negative turns. The first approach is to fit the cooperative McGhee-Von Hippel binding model simultaneously to the change in contour length (Figure 2d,g) and to the shifts of the center positions of the extension-rotation curves (Figure 3b,e), with the dissociation constant *K*_D_, the binding site size *n*, cooperativity parameter *ω*, and unwinding angle per molecule Δ*θ* as fitting parameters (Equations 1and 4). We find excellent agreement of the binding curves (Figure 2d,g and Figure 3b,e) for both SxO and SxG and fitted unwinding angles of Δ*θ* = 21.1° ± 0.1° for SxO and Δ*θ* = 20.5° ± 0.1° for SxG. Alternatively, we directly analyzed the shift in the center of the rotation curves as function of measured contour length and find a linear relationship (Figure 3g,h). Using Equations 2 and 3, we expressed *ΔTw* as a function of *L*_C_ with a slope of Δ*θ*/Δ*z*. Again, assuming a length increase of 0.34 nm per intercalation event, the fitted linear slopes give an unwinding angle per molecule of Δ*θ* = 21.5° ± 0.6° for SxO and Δ*θ* = 20.1° ± 0.5° for SxG (errors obtained by the covariance matrix of the fit), in excellent agreement with the result from the fits of the McGhee-von Hippel model. The observed unwinding angle per molecule for both SxO and SxG falls in the middle of the range of values observed for other monointercalators ^35, 37, 39, 44^ (Figure 3i).

### Intercalation of SxO and SxG has only a small effect on plectoneme size

The slopes of the extension-rotation curves in the linear, plectonemic regime (Supplementary Figure S10) provide a measure for the size of the plectonemes ^51, 52, 53^. Throughout the entire SxO concentration range probed, the slopes of the plectonemic regime remain essentially constant, within error (Figure 3c). For SxG, the slopes of the plectonemic regime remain within error until a concentration of 1 μM, and then show a small, but consistent decrease (Figure 3f). Together, the results suggest that for both SYTOX dyes the size of plectonemes is not affected by intercalation in the range typically used for visualization ^17, 21, 22, 23^ (≤ 1 µM), which is advantageous in measurements that visualize and probe DNA supercoils and loops and their interactions and processing.

### Evidence for torque-dependent binding

We observe that in the presence of SxO or SxG, the rotation curves are much broader than in the absence of dye and that the maximum extension does not occur at the center of the curve, as is the case for bare DNA, but instead just before the buckling point for the formation of negative plectonemes (Figure 3a,d). The length increase indicates increased intercalation when the DNA is underwound and can be explained by torque-dependent binding ^25, 28, 54^. A negative torque applied to the right-handed DNA helix favors intercalation, by Le Chatelier’s principle, as intercalation causes the DNA to unwind. Specifically, an applied torque *Γ* changes the net free energy needed for SxO or SxG to intercalate by the work done as *Γ*·Δ*θ*, where Δ*θ* is the unwinding angle upon intercalation ^25, 28, 54, 55^. Increased lengthening caused by intercalation upon applying negative turns (and shortening at positive turns, all relatively to the torsional relaxed molecule at the given dye concentration) explains the apparent slope in between the buckling points and contributes to the broadening of the rotation curves.

### Intercalation of SxO and SxG stabilizes B-form double-stranded DNA and suppresses torque induced melting and P-DNA formation

To further characterize the effect of SxO and SxG intercalation on DNA conformations, we measured rotation-extension curves at elevated forces (> 1 pN), both in PBS buffer and in the presence of a high concentration (20 μM) of SxO or SxG (Figure 4 and Supplementary Figure S11). For bare DNA at forces > 1 pN, buckling and the formation of pletonemic supercoils is suppressed upon underwinding, due to torque induced melting ^56, 57, 58^. Above 5 pN, DNA no longer buckles even when overwound, due to the formation of an alternate structure called P-DNA ^59, 60^ (Supplementary Figure S12). At high concentrations of SxO or SxG, buckling occurs at both negative and positive turns for all stretching forces tested (0.5–7 pN; Figure 4 and Supplementary Figure S11), suggesting that both torque-induced melting and P-DNA formation are suppressed. The findings indicate that intercalation of SxO and SxG into the double-stranded B-form of DNA is preferred relative to torque-induced melted or P-form DNA, which in turn suppressed the formation of these alternative DNA structures in the presence of high concentration of SxO or SxG. The observed stabilization against torque-induced melting is in line with the increase in the melting temperature *T*_m_ of DNA in the presence of 20 μM SxO shown previously ^61^ and consistent with similar effects seen for other intercalators ^35^.

**Figure 4.**
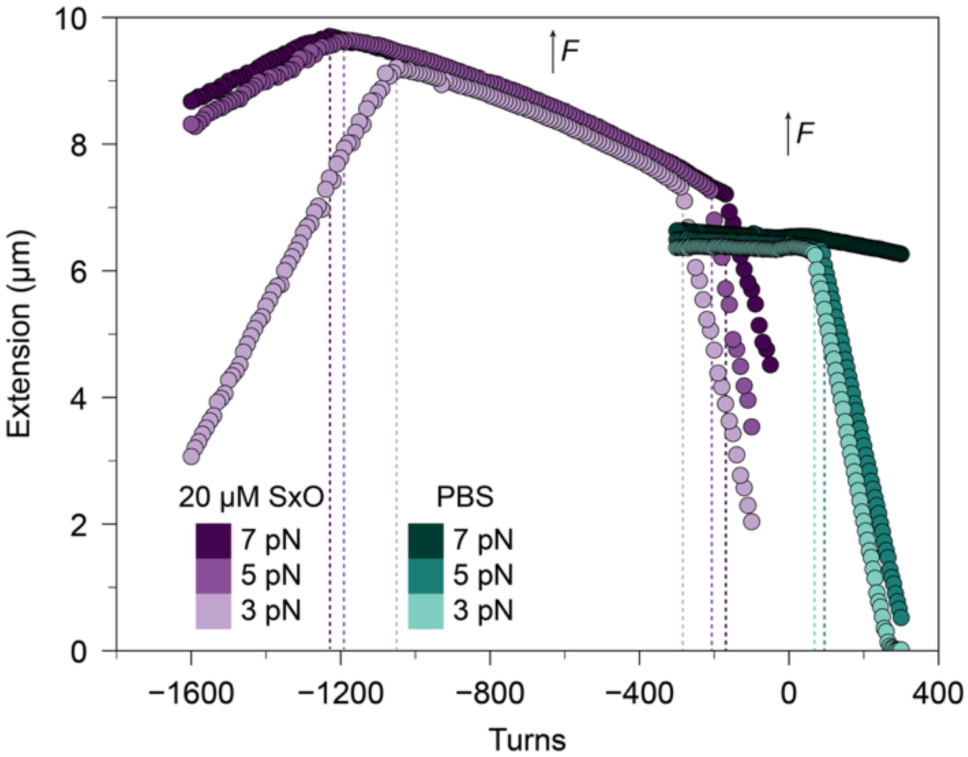
Rotation-extension measurements at high forces show a stabilization of B-form, double-stranded DNA by SxO. A) Rotation-extension curves for 20.6 kbp DNA in PBS (teal shades) and 20 μM SxO (purple shades) at stretching forces of (light to dark colors) 3, 5 and 7 pN. The presence of SxO significantly affects the positions and shape of the rotation-extension curves. Vertical dashed lines indicate the buckling points, i.e. the onset of plectoneme formation, for positive supercoils in PBS and for both positive and negative supercoils in 20 μM SxO. High force rotation-extension curves of DNA in the presence of SxG are shown in Supplementary Figure S11.

### SxO and SxG intercalation shifts their absorbance spectra to longer wavelengths

To correlate the results from MT measurements to their optical spectroscopic properties, we measured the absorbance of both SxO and SxG in the absence and presence of defined concentrations of DNA (Materials and Methods and Figure 5). The absorbance spectrum of SxO in the absence of DNA is characterized by a main absorption band at a wavelength of *λ* ≈ 529 nm and a minor band around ≈ 300 nm (Figure 5 and Supplementary Figure S13). Similarly, for SxG in the absence of DNA, the spectrum features a main absorption band at *λ* ≈ 500 nm and two minor bands around ≈ 300 nm and ≈ 360 nm (Figure 5 and Supplementary Figure S14). Upon the addition of DNA, the absorbance spectra of SxO and SxG exhibit bathochromicity, i.e. a shift to longer wavelengths. For SxG the shift is small, from *λ*_free_ ≈ 499 nm to *λ*_bound_ ≈ 504 nm (Figure 5d). For SxO, the spectral changes upon addition of DNA are much more pronounced and the main absorbance peak shifts from 529 nm to 548 nm (Figure 5a). The bathochromic shifts upon binding to double-stranded DNA are similar to what has been observed for other intercalators ^62^.

**Figure 5.**
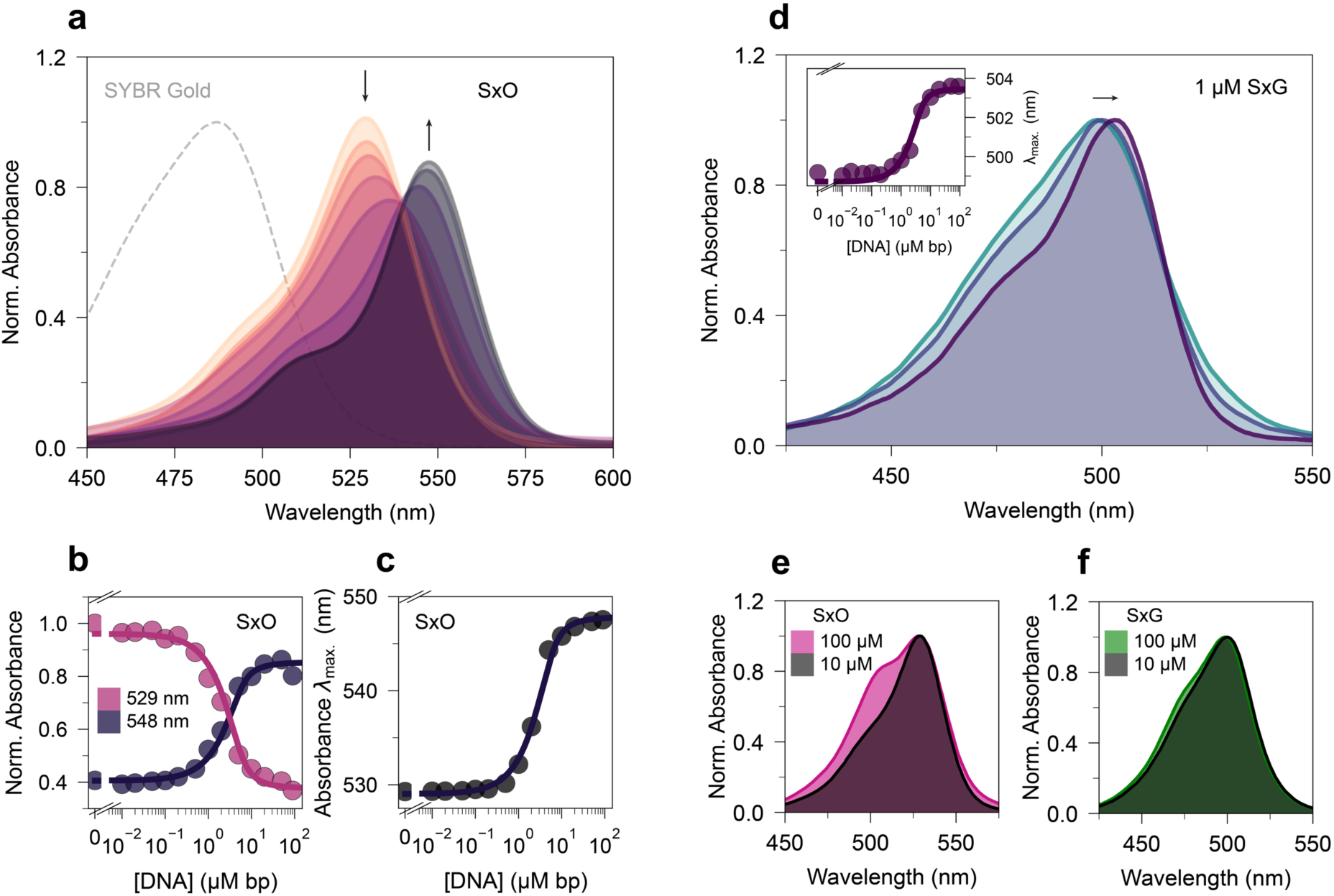
Absorbance spectra of SxO and SxG show systematic changes upon intercalation. a). Absorbance spectra of a 1.7 μM SxO solution in PBS upon titration with λ-DNA. The base pair concentrations shown are (light to dark) 0, 0.1, 0.5, 1, 2, 5, 20, 50 μM. Data are normalized to the maximum absorbance in absence of DNA. The dashed line is the absorbance spectrum of SYBR Gold ^39^ for comparison. b) Absorbance intensity at 529 nm and 548 nm of 1.7 μM SxO as function of DNA concentration. The solid line is a fit to the finite concentration McGhee-Von Hippel binding model (Methods) with the absorbance values *A*_free_ and *A*_bound_ as fitting parameters. We find values of 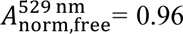 and 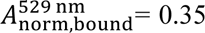 for *λ* = 529 nm and values of 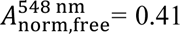 and 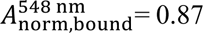 for *λ* = 548 nm from the fit. Data are from spectra in panel a. c) Wavelength of the absorbance maxima as function of DNA concentration. The solid line is a fit to the finite concentration McGhee-Von Hippel binding model with the wavelengths *λ*_max,free_ and *λ*_max,bound_ as fitting parameters. We find values of *λ*_max,free_=529 nm and *λ*_max,bound_=548 nm from the fit. Data are from spectra in panel a. d) Absorbance spectra of a 1 μM SxG solution in PBS upon titration with λ-DNA. The base pair concentrations are (light to dark) 0, 2 and 90 μM. Data are normalized to the maximum of each spectrum. The inset shows the wavelength of maximum absorbance as function of DNA concentration. The solid line is a fit of the finite concentration McGhee-Von Hippel binding model with *λ*_max,free_ =499 nm and *λ*_max,bound_ =503 nm determined from the fit. e) Absorbance spectra of SxO at high concentrations in PBS. Spectral deformation suggests intermolecular dye-dye interactions. f) Absorbance spectra of SxG at high concentratios in PBS.

For both dyes, the shift in wavelength with increasing DNA concentration can be well-described by a simple model (Equation 6) that used the finite-concentration McGhee-von Hippel model to compute the amount of intercalation and the absorption wavelengths of the free and bound dye as fitting parameters (Figure 5c and d, inset). For SxO, we find excellent agreement when using the binding parameters obtained from the MT measurements (Figure 2d). For SxG, the binding parameters from MT measurements (Figure 2g) give a reasonably albeit not fully accurate description (Supplementary Figure S15) and we, therefore, use binding parameters obtained using fluorescence spectroscopy (see below).

For SxO, we observe, in addition to the shift in wavelength, a decrease of ≈ 10% in absorbance intensity upon intercalation. The changes in absorbance intensity, evaluated at the wavelength of the maxima of the free or bound SxO (Figure 5b), respectively, can similarly be described as a linear superposition of free and bound dye contributions (Equation 5), using the binding parameters determined from the MT measurements and the absorbances of the free and bound dye as fitting parameters. The value for 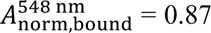 suggests a decrease of 13% in absorbance intensity upon intercalation, in good agreement with the ratio of molar absorption coefficients of the DNA-dye complex and the free dye 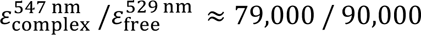 (Supplementary Figure S13b) ^5^. Overall, the agreement between the binding parameters found using MT and the spectral shifts of the absorbance strongly suggests that the same intercalative binding mode is observed through single-molecule mechanics and bulk absorbance spectroscopy.

### Intermolecular interactions at high SxO and SxG concentrations

To investigate the behaviour of the SYTOX dyes and possible interdye interactions at high concentrations (> 10 µM), we measured the absorbance of SxO and SxG at elevated concentrations (Figure 5e,f). For SxO, we observe changes in the spectrum above 10 <u>µ</u>M that are characteristic of intermolecular interaction of cyanine dyes ^63^. Specifically, we observe more pronounced absorbance around 500 nm, blue shifted compared to the maximum at low dye concentration, which is suggestive of H-dimerization (Figure 5e). H-dimers (or H-aggregates) are side-by-side configurations of dye molecules with a blue-shifted absorbance compared to the monomer. Addition of DMSO reduces the spectral features indicative of interdye interactions (Supplementary Figure S16), likely due to creating a less polar environment with a higher solubility for the SYTOX dyes. For SxG similar spectral changes are detectable at high concentrations, but they are much less pronounced compared to SxO (Figure 5f), likely because of larger electrostatic repulsion due to SxG’s triple negative charge.

### SxO and SxG intercalation leads to a large increase in fluorescence

SxO and SxG are widely used as fluorescent DNA stains. To characterize their fluorescence in the absence and presence of DNA, we recorded fluorescence emission spectra (Figure 6). We find that adding DNA to a fixed concentration of SxO or SxG causes a large increase in fluorescence intensity together with a bathochromic shift in the fluorescence wavelengths (Figure 6a-d). The maximum fluorescence emission intensity of the DNA-dye complex is at *λ*=570 nm for SxO and at *λ*=523 nm for SxG, in excellent agreement with the wavelengths reported by the manufacturer (570 nm and 523 nm). In the absence of DNA, we observe a fluorescence emission maximum at *λ*=556 nm for SxO (Figure 6e) and at ≈520 nm for SxG (Supplementary Figure S17). While the solvent environment has only relatively modest effects on the absorbance and emission wavelengths of SxG, for SxO the absorbance and fluorescence spectra shift systematically to longer wavelengths going from PBS, to DMSO, to the presence of DNA (Figure 6e). The observation of negative solvatochromism for SxO, i.e. a shift to longer wavelengths upon decrease of the polarity of the environment is in line with previous reports for cyanine dyes ^64, 65^. Excitation spectra of SxO (collected at *λ*_em_ = 600 nm) resemble the observations for the absorbance and show a systematic shift in excitation maximum upon binding to DNA (Supplementary Figure S18).

**Figure 6.**
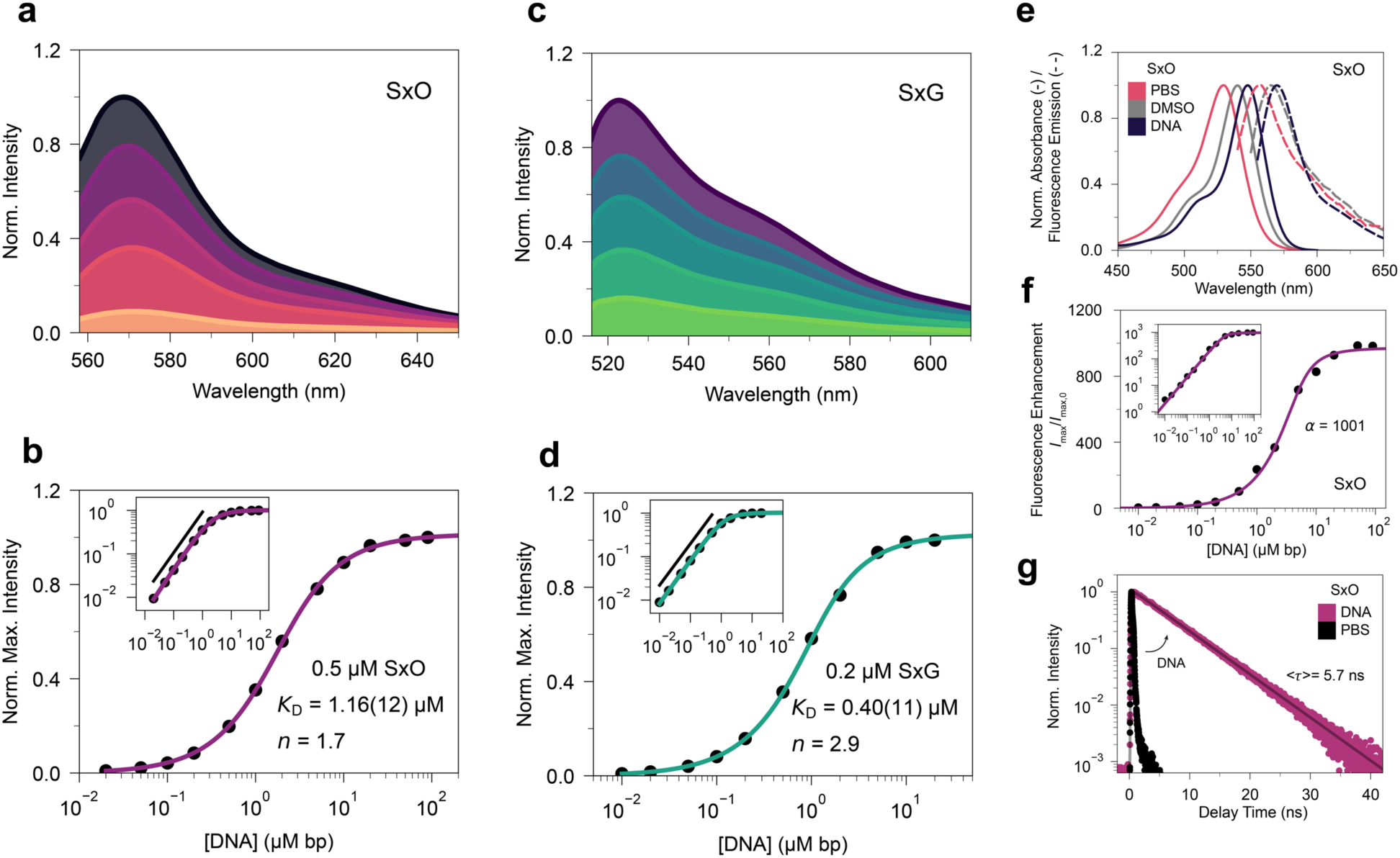
Fluorescence emission spectra show a large increase of SxO and SxG fluorescence upon intercalation. a) Fluorescence emission spectra of 0.5 μM SxO in PBS with increasing concentrations of λ-DNA. DNA concentrations are (light to dark) 0.2, 1, 2, 5, 50 μM·bp. b) Fluorescence enhancement of 0.5 μM SxO solution in PBS upon addition of DNA. The solid line is a fit of the finite concentration non-cooperative McGhee-von Hippel binding model with parameters given in the legend. Samples were excited at 504 nm. Data are normalized to the signal at the highest DNA concentration. The inset shows the same data on a log-log scale. The black solid line is linear fit to the data ≤ 1 μM·bp, shown offset for clarity, with a fitted slope of 0.94, suggesting a (close to) linear response to increasing DNA concentration. c) Fluorescence emission spectra of 0.2 μM SxG in PBS upon titration of λ-DNA. DNA concentrations are (light to dark) 0.2, 0.5, 1, 2, 30 μM·bp. d) Fluorescence enhancement data similar to panel b, but for 0.2 μM SxG. The black line is fitted to the data ≤ 0.5 μM bp, shown offset for clarity, and gives a slope of 0.95. e) Absorbance and fluorescence emission spectra of SxO in PBS, DMSO and in the presence of DNA. f) Fluorescence enhancement of 1.7 μM SxO solution in PBS upon the addition of λ-DNA, relative to the absence of DNA. The solid line is a fit of the finite concentration McGhee-von Hippel binding model using the binding parameters found from MT measurements treating the fluorescence enhancement *α* as fitting parameter. We find a value of *α* = 1001 from the fit. Samples were excited at 547 nm. g) Time-Correlated Single-Photon Counting (TCSPC) measurements of 2 μM SxO solution in PBS and 0.2 μM SxO in the presence of 2 μM·bp DNA. The solid line is a mono-exponential decay fit to the data with a fluorescence lifetime <*τ*> = 5.7 ns. The fluorescence lifetime of free SxO is faster than the reliable time resolution of our instrument, < 0.3 ns.

We observe a large and systematic increase in the fluorescence emission intensity with rising DNA concentrations for SxO and SxG (Figure 6b,d). For an excess of dye, the increase in fluorescence is linear with the DNA concentration (Figure 6b,d insets), which is a basis for the quantification of nucleic acids. We find that the fluorescence emission enhancement as a function of DNA concentration is well described by the noncooperative McGhee-Von Hippel binding model at relatively low binding densities (Figure 6b,d, solid lines), with an apparent binding affinity of *K*_D_ = 1.2 μM and binding site size of *n* = 1.7 for SxO and *K*_D_ = 0.43 μM and binding site size of *n* = 2.8 for SxG. The binding affinities found are similar but slightly higher than values for the anticooperative model using MT, likely due to absence of a cooperativity factor < 1. We note that for high SxO concentrations (> 2 µM), the fluorescence intensity decreases again with increasing dye concentration (Supplementary Figure S19), so this regime should be avoided for DNA quantification, as previously observed for SYBR Gold ^39^.

To quantify the fluorescence emission enhancement of SxO upon binding to DNA, i.e. the increase in intensity with respect to the initial maximum intensity (*I*_max_/*I*_max,0_), we recorded its fluorescence intensity relative the dye in absence of DNA (Figure 6f). The data suggest a ∼1000 fold increase in maximum fluorescence emission of SxO upon DNA intercalation, which is similar to other intercalating nucleic acid stains ^11, 62, 66, 67^. We compute the free and bound SxO concentration from the finite concentration cooperative McGhee-von Hippel model using the binding parameters *K*_D_, *n* and *ω* determined from the MT measurements and find an excellent fit to the fluorescence binding curve, using the fluorescence enhancement as a free fitting parameter. The excellent agreement between the fluorescence enhancement and the binding parameters determined from mechanical manipulation further confirms that intercalation of the dye is responsible for its increased quantum yield and that both modalities quantify intercalation.

### Fluorescence lifetime measurements show an increase in fluorescence lifetime upon intercalation

To further investigate the fluorescence properties of SxO, we turned to time-correlated single-photon counting (TCSPC) measurements to determine its fluorescence lifetime (Materials and Methods and Figure 6g). The fluorescence lifetime of free SxO is faster than the reliable time resolution of our instrument, < 0.3 ns. In contrast, in complex with DNA the lifetime is much longer, *τ* = 5.7 ns (0.2 μM SxO; dye/DNA = 1 / 10 bp) and is well described by a single exponential decay in the presence of an excess of DNA, close to the structurally similar SYBR Gold ^39^ with a fluorescence lifetime of *τ* = 6.1 ns and similar to other intercalating nucleic acid stains ^62, 66, 67^. A single exponential decay is indicative of one dominant emissive state and a homogeneous environment, suggesting an overall uniform interaction with the DNA bases. An increase in lifetime together with the large increase in emission intensity agrees with conformational restriction upon intercalation of the dye, as proposed for other cyanine dyes.

### Quantum chemical calculations relate structural to spectroscopic properties

We perform quantum chemical calculations (see Methods) to investigate the SxO and SxG structures and to relate them to spectroscopic properties. The optimized ground state structures show planar fluorophore geometries for SxO and SYBR Gold and mostly planar geometries with a small tilt for SxG and PicoGreen, with side chains that protrude from the plane of the quinoline and aza-benzoxazole or benzothiazole moieties, respectively (Figure 7 and Supplementary Figure S20). For all four molecules and for both water and DMSO solvent parameters the excitation from the ground to the first excited state is the brightest of the first ten excited states (Supplementary Tables S2 and S3). This excitation is always dominated by a HOMO to LUMO transition with a weight of more than 88%.

**Figure 7.**
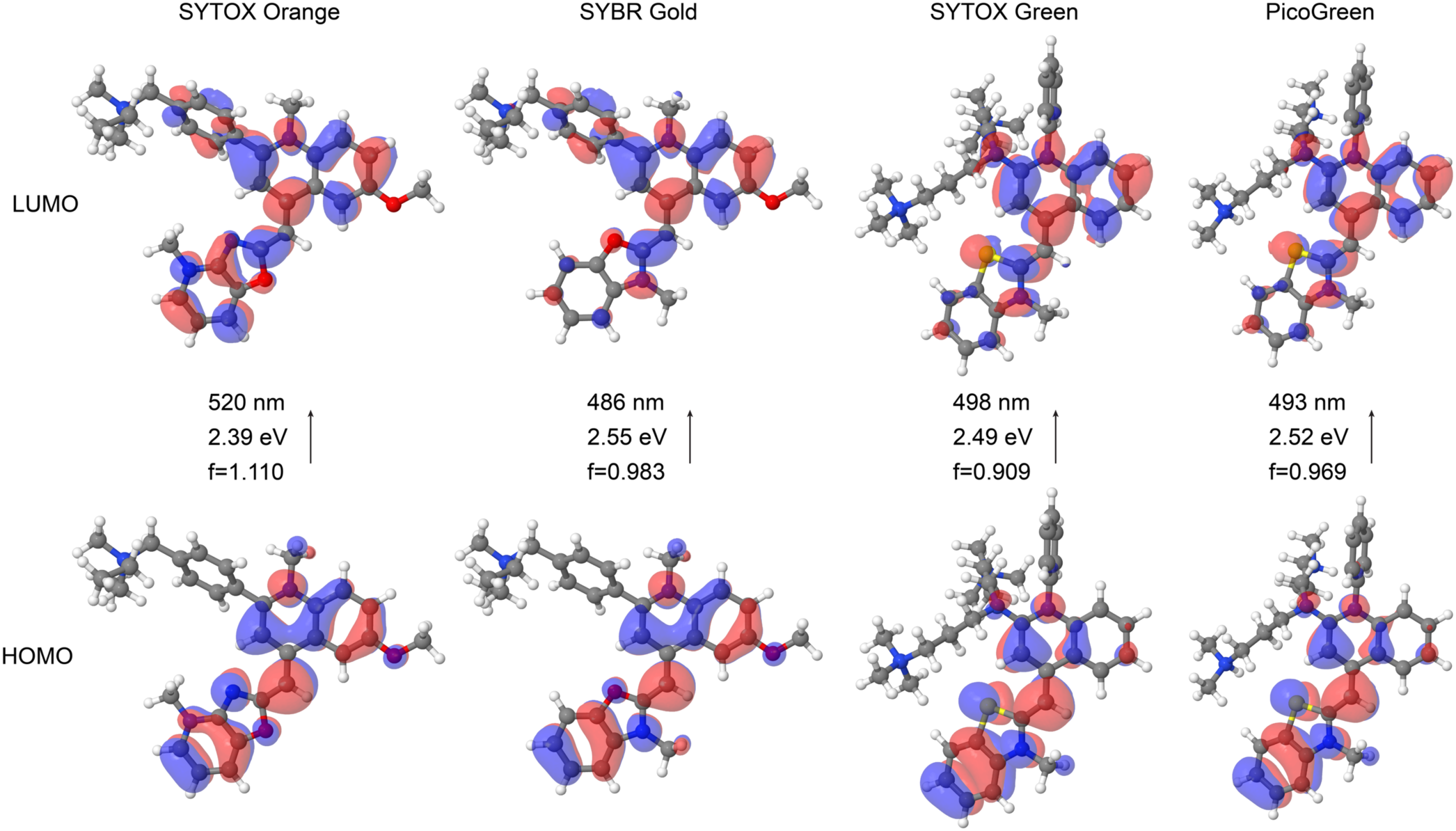
Quantum chemical calculations predict ground state structures and absorption wavelengths. Structures for SxO, SYBR Gold, SxG, and PicoGreen were optimized at the RI-CC2/cc-pVDZ level of theory and the subsequent excited state calculations with RI-CC2/aug-cc-pVDZ employing the COSMO solvation model with parameters representing water as solvent were used for orbital visualization (Materials and Methods). The highest occupied molecular orbitals (HOMO; bottom) and lowest unoccupied molecular orbitals (LUMO; top) are shown. The excitation energies form the ground to the first excited state, the corresponding wavelengths, and the oscillator strengths are indicated for each dye. For each of the four molecules, this excitation is dominated by a transition from HOMO to LUMO. The orbitals and corresponding predicted absorption wavelengths are rather similar for SxG and PicoGreen, in agreement with experimental data and consistent with the very similar core structure. For SxO and SYBR Gold, the orbitals show clear differences, in particular for the LUMO, and the predicted absorption wavelength is considerably red shifted for SxO. The results from calculation using DMSO solvent parameters are shown for comparison in Supplementary Figure S20.

For SxG and PicoGreen, the molecular orbitals are virtually identical and the coupled-cluster calculations predict similar absorption wavelengths (Supplementary Figure S21 and Supplementary Tables S2 and S3), in good agreement with the experimental observation of very similar absorbance maxima for SxG and PicoGreen, which differ by only a few nm (Table 2).

**Table 1.** Properties of SYTOX Orange and SYTOX Green. Molar absorptivity coefficients were recorded in PBS buffer, at a wavelength of *λ*_max_ = 529 nm for SxO and at *λ*_max_ = 500 nm for SxG.

| Dye | Molar Absorptivity Coefficient $\epsilon$ ($10^4 \text{ M}^{-1} \text{ cm}^{-1}$) | Chemical Formula | Molecular Weight (u) | Counterion |
| --- | --- | --- | --- | --- |
| SYTOX Orange | 9.0 | $\text{C}_{31}\text{H}_{36}\text{N}_4\text{O}_2^{2+}$ | 496.65 | 2 $\text{I}^-$ |
| SYTOX Green | 7.4 | $\text{C}_{36}\text{H}_{48}\text{N}_5\text{S}^{3+}$ | 582.87 | 3 $\text{I}^-$ |

**Table 2.** Absorption, excitation, and emission wavelengths for SYTOX Orange, SYTOX Green, SYBR Gold and PicoGreen. Data for SxO and SxG were obtained in this work. Data for SYBR Gold are from ^39^ and data for PicoGreen from ^62^.

| Dye | Solvent/Complex | Abs/Exc/Em (nm) | Stokes shift (nm)/(cm<sup>-1</sup>) |
| --- | --- | --- | --- |
| SYTOX Orange | PBS | 529/529/556 | 27/918 |
|  | DNA | 548/548/570 | 22/704 |
|  | DMSO | 540/-/565 | 25/819 |
| SYTOX Green | PBS | 500/-/≈520 | ≈20/≈770 |
|  | DNA | 504/505/524 | 20/757 |
|  | DMSO | 504/-/- | - |
| SYBR Gold | PBS | 486/≈485/≈550 | 64/2394 |
|  | DNA | 494/496/543 | 49/1827 |
| PicoGreen | PBS | 498/-/528 | 30/1141 |
|  | DNA | 501/-/522 | 21/802 |

Experimentally, the absorbance maxima of free and bound SxO exhibit a large bathochromic shift (i.e. to longer wavelength) compared to the structurally very similar SYBR Gold (by 42 nm and 54 nm, respectively; Table 2), indicating that the incorporation of an additional nitrogen atom in the benzoxazole moiety results in a significant spectral shift (Figure 5a). A similar bathochromic shift of the absorption of SxO compared to SYBR Gold is reproduced in our excited state calculations. The frontier orbitals from these calculations show relatively large differences between SYBR Gold and SxO. The difference is particularly notable for the LUMO, where there is a more pronounced contribution from the aza-benzoxazole moiety in SxO than from the benzoxazole in SYBR Gold. This difference reflects a larger conjugated system and is the origin of the bathochromic shift in absorption for SxO.

## DISCUSSION

We resolved the molecular structure of SxO and SxG and found SxO to be an aza-benzoxazole containing cyanine dye, containing an additional nitrogen in its core and otherwise identical to SYBR Gold. We found SxG to be closely related to PicoGreen with two additional N-methyl groups. Absorbance and emission spectroscopy show that the additional nitrogen atom of SxO results in a significant bathochromic shift compared to the closely related SYBR Gold, in particular in the absorption spectrum. In contrast, the spectra of SxG are very similar to the related dye PicoGreen, indicating that the side chain methyl substitutions have only minor effects on the optical properties.

We employed single-molecule MT measurement to probe DNA under controlled forces and twist in the absence and presence of different concentrations of SxO and SxG. The MT data reveal a lengthening of DNA by approximately 50% at high dye concentrations and unwinding by Δ*θ* =\ 21.1 ± 0.1° per SxO molecule and Δ*θ* = 20.5 ± 0.1° per SxG molecule, characteristic of intercalation.

Fits of the anti-cooperative McGhee-von Hippel binding model describe the data where no clear saturation occurs in the measured concentration range. A cooperativity factor < 1 has previously been reported for ethidium bromide ^44^. The binding parameters found through single-molecule mechanics are in excellent agreement with bulk absorbance and fluorescence spectroscopy, confirming that the single, intercalative binding mode is responsible for the binding behavior seen in the optical spectroscopy and by mechanical manipulation. Changes in the bending persistence length remain within error over a large range of SxO and SxG concentrations and for dye concentrations < 1 µM the bending stiffness changes by < 10%, relative to DNA in the absence of dye. Similarly, the slopes in the plectonemic regime, proportional to the size of plectonemes, remain constant in the concentration range typically used for staining of DNA (< 1 μM SxO). Small or negligible changes in DNA mechanics are advantageous for dyes used to visualize and track DNA loops and supercoiling, where the DNA properties should ideally remain close to the behavior in the absence of dye. The small effect on mechanical properties combined with the ∼1000-fold increase in fluorescence emission upon intercalation for SxO and even higher for SxG make them excellent choices for supercoil and plectoneme visualization assays. Together, our data provide a comprehensive characterization of the popular DNA stains SxO and SxG using several complementary methods. Our findings provide several key parameters for modeling DNA in the presence of both dyes and serve as a guideline for future assay and fluorophore development.

## METHODS

### Chemicals and preparation of dilution series

SYTOX Orange and SYTOX Green were obtained from Invitrogen as a stock solution of 5 mM in DMSO (SYTOX Orange Nucleic Acid Stain, cat# S11368, Invitrogen; SYTOX Green Nucleic Acid Stain, cat# S7020, Invitrogen), or alternatively 250 μM in DMSO for comparison (SYTOX Orange Dead Cell Stain; cat# S34861, Invitrogen) (Supplementary Figure S5). The DMSO concentrations in the buffer solutions containing the dye were ≤ 2% (v/v), or ≤ 20% (v/v) when using the 250 µM stock for comparison.

### Mass Spectrometry

Mass spectrometric analyses were performed using a timsTOF instrument (Bruker) operated in direct infusion mode. The dyes, supplied in DMSO, were diluted in pure methanol and introduced into the mass spectrometer via a syringe pump. Electrospray ionization (ESI) was applied in both positive and negative ion modes. Nitrogen was used as the drying gas at a flow rate of 8 L min⁻¹, a pressure of 1.8 bar, and a temperature of 200 °C. Mass spectra and fragment ions were acquired in auto MS/MS mode over an *m/z* range of 50–1300, with a resolving power of 40,000 and a mass accuracy of approximately 0.3 ppm. Data acquisition was carried out using Bruker Compass HyStar 5.1 (v5.1.8.1) and otofControl 4.1 (v6.2), and data analysis was performed with DataAnalysis 5.0 (vSR1). The isotope distribution was calculated using ChemDraw 26.

### NMR Spectroscopy

NMR samples of SYTOX Green and SYTOX Orange were prepared by lyophilizing the commercially available stock solutions (250 μL, 5 mM in DMSO) and dissolving the dye in 180 μL of DMSO-d_6_ (99.96% D, Eurisotope). For measurements below 290 K 25 vol-% of D_2_O were added to lower the freezing point ^68^. NMR measurements were performed at spectrometers with ^1^H resonance frequencies of 500, 600, 800, 900, 950 and 1200 MHz (Bruker Biospin) equipped with inverse triple or quadruple resonance probes (mostly He-cooled cryoprobes). Spectra were generally recorded at 298 K, but also at various temperatures between 253 K and 358K (for SYTOX Orange). Sample temperatures were calibrated with a 99.8% methanol-d_4_ reference sample (Bruker).

For the complete assignment of the ^1^H, ^13^C and ^15^N signals a set of 2D experiments was used (DQF-COSY, TOCSY, ^1^H,^13^C-HSQC, ^1^H,^13^C-HMBC, ^1^H,^15^N-HSQC, ^1^H,^15^N-HMBC), in addition to 1D, ^1^H and ^13^C spectra. Sample impurities were identified from 2D DOSY spectra, relying on differing diffusion constants of dye and impurities. 2D NOESY and ROESY spectra were used to collect NOE data for confirming assignments and the connectivity of core and tail fragments. All 1D and 2D spectra were processed and evaluated using the TOPSPIN software package (versions 3.5pl7 and 3.8.0, Bruker Biospin).

### DNA construct and flow cell preparation for magnetic tweezers measurements

Our magnetic tweezers measurements used a 20.6 kbp (46% GC content) DNA construct, based on the λ-phage DNA sequence and prepared as described previously ^35, 69^. The DNA construct has (≈600 bp) PCR-generated handles with multiple biotin or digoxigenin moieties ligated to the unmodified central sequence of the DNA. Streptavidin coated MyOne superparamagnetic beads (Dynabeads MyOne Streptavidin C1, cat# 65001, Thermo Fisher Scientific) with 1 μm diameter are attached to the DNA constructs through multiple bonds to the biotin handles by incubating 2 μL of 0.2 ng μL^-1^ DNA stock solution with 2 μL of the magnetic beads in 100 μL 1× PBS (10 mM phosphate buffer, pH 7.4, with 138 mM NaCl and 2.7 mM KCl; cat# P3813, Sigma-Aldrich) for 5 minutes.

Flow cells are prepared using two microscope coverslips (24x60 mm #1, Menzel) (Supplementary Figure S6). The bottom coverslip was functionalized with 3-Glycidoxypropyltrimethoxysilane (cat# 10791101, Fisher Scientific) and incubated with 100 μL of a 5000-fold diluted stock solution of 1 μm diameter polystyrene beads (Polybead Microspheres 1.00 μm, Polysciences) in absolute ethanol used as reference beads. To allow for liquid exchange, the top coverslip was prepared with two openings with a 1 mm radius. The flow cell is assembled by stacking the two coverslips using a layer of Parafilm (Carl Roth), with a central ∼ 25 μL channel cut out, and sealed on a hotplate (90 °C) for 20 seconds. This channel is connected to the two openings of the top coverslip functioning as an inlet and an outlet for fluid exchange.

After flow cell assembly, 100 μL of 100 μg mL^-1^ anti-digoxigenin (cat# 11333089001, Roche) in 1× PBS is introduced and incubated for at least one hour. The flow cell is passivated using 300 μL of 50 mg mL^-1^ bovine serum albumin (cat# A2153, Sigma Aldrich) in 1× PBS for at least one hour to counter non-specific interactions. The bead-coupled DNA constructs are introduced in the flow cell and allowed to bind via multiple digoxigenin:anti-digoxigenin interactions on the bottom coverslip for 30 minutes.

To test for equilibration and nonspecific binding of the dye to the flow cell, rotation-extension curves were recorded in PBS and subsequently after flushing with various volumes of 0.1 μM SxO (Supplementary Figure S7). The degree of DNA unwinding was monitored three or four times in the span of approximately 30 min to test system equilibration. The measured DNA unwinding after a flushing volume of 600 μL remained within ±3% after approximately 25 min or after repeated flushing of an additional 600 μL of dye solution.

### Magnetic tweezers setup

We use a custom-built MT setup that was described previously ^40, 70^, employing two 5 × 5 × 5 mm^3^ permanent magnets (W-05-N50-G; Supermagnete) with a 1 mm gap placed in a vertical configuration ^30^ mounted on a DC translational motor (M-126.PD2, PI) to control the distance between magnets and the flow cell, together with a rotational motor (C-150.PD, PI) to control the rotation of the magnets. For illumination, we used an LED (69647; Lumitronix LED Technik) and for imaging a 40× oil-immersion objective (UPLFLN 40×; Olympus) projecting to a CMOS sensor camera with 4096 × 3072 pixels (12M Falcon2; Tele-dyne DALSA) to image a field of view of 400 × 300 μm^2^. Images recorded at 58 Hz were transferred to a frame grabber (PCIe 1433; National Instruments) and 3D bead positions were tracked in real time using custom-written tracking software (Labview, National Instruments) ^71, 72^. A z-position look-up table was created by controlling the objective position using a piezo stage (PifocP726. 1CD, PI Physikinstrumente) over a range of 20 μm, with a step size of 100 nm. Labview routines described previously were used for set up control and bead tracking ^72^.

### Magnetic tweezers measurements

Prior to the introducing dyes, the DNA tethered beads are subjected to force and torque to detect the binding to multiple tethers and to test for torsional constraint. First, negative turns are applied under high applied force (*F* = 5 pN). At high forces, rather than buckling to form plectonemes, melting of DNA occurs upon applying negative turns ^56^ (corresponding to underwinding of the DNA), preventing a decrease in extension (Figure 3A). However, in the case of multiple tethers attached to the same bead, a decrease in extension is still observed as the tethers braid upon rotation of the magnetic bead. Beads bound to multiple tethers are not analyzed further. To test for torsional constraint of the DNA tethers, positive turns are applied at low tension (*F* = 0.5 pN). In the case of torsionally constrained DNA, the tethers will overwind and form plectonemes, consequently decreasing the extension. In contrast, torsionally unconstrained tethers will not overwind and no plectonemes are formed.

First, rotation-extension measurements are performed on torsionally constrained tethers by determining the extension while varying the relative turns of the magnet. The linking number is changed in steps of five turns, after which the extension is measured for 10 s and averaged. To determine the initial orientation of the tethers, a rotation curve is measured in 1×PBS. While applying a force of 0.5 pN, the magnets are rotated positively or negatively and the extension is monitored. From this, the number of turns to achieve fully relaxed tethers (corresponding to the center of the rotation curve) is determined. Second, force-extension curves are recorded by measuring the tether extension upon setting the magnet to 12 different positions above the flow cell, corresponding to forces between 0.05 - 5 pN. Forces were calibrated using the bare DNA tethers by determining the transverse fluctuations using the equipartition theorem ^29, 33^.

Next, using a peristaltic pump 600 - 1000 μL (corresponding to > 25 flow cell volumes) of SxO or SxG solution in 1×PBS at defined concentration are introduced to the flow cell, in order from low to high concentration. A force of 2 pN is applied while flushing to constrain movement of the bead, in particular to ensure no change in bead rotation during fluid extension. Next, the rotation-extension measurement at low force (0.5 pN) is repeated. The linking number is scanned in steps of five turns for low concentrations and in steps of 10 turns for 0.2 μM and higher, because of the large number of turns required. Afterwards, the force-extension relation is measured. This procedure is applied subsequently to all SxO and SxG dilutions.

### Effects of the presence of DMSO

Upon supplementation of the buffer with dye, a small amount of DMSO is added simultaneously, as it is the solvent of the stock solution, which might affect the binding properties of the dye and mechanics of the DNA ^40^. We tested the effect of DMSO by monitoring the unwinding of the DNA upon intercalation through MT rotation-extension curves, for different amounts of DMSO background (Supplementary Figure S5). We find that as the concentration of DMSO approaches 10% (v/v), the degree of binding starts to deviate from the binding observed for a much lower DMSO concentration of < 0.5% (v/v). Therefore, for the measurements reported in the main text we kept the DMSO concentration to ≤ 2% DMSO.

### Absorbance, excitation, and emission spectra

Absorbance and fluorescence excitation and emission spectra of SxO and SxG were collected in PBS, unless stated otherwise, and corrected for background signal. DNA titrations were prepared from a stock of λ-DNA (50% GC content; 48.5 kbp) (Lambda DNA, 0.3 μg μL^-1^, 10 mM Tris-HCl (pH 7.6) and 1 mM EDTA; cat# SD0011, Thermo Scientific). The samples were protected from light and left to equilibrate for at least 10 min. We employed λ-DNA, instead of the functionalized DNA construct used in MT measurements, since the functionalized DNA is difficult and wasteful to produce in sufficient quantities for spectroscopy and since the labels used for attachment in the MT might interfere in the optical spectroscopic measurements.

SxO absorbance spectra were collected on a PerkinElmer LAMBDA 365 UV/Vis Spectrophotometer. SxO fluorescence excitation and emission spectra were collected on a PerkinElmer FL6500 Fluorescence Spectrophotometer. All SxO samples were measured in plastic UV-transparent cuvettes with a pathlength of 10 mm (BRAND UV cuvettes, cat# 759125, Sigma-Aldrich). Additionaly, SxO fluorescence spectra in the presence of λ-DNA were collected on a Tecan INFINITE M1000 PRO in a 96-Well Glass Bottom Matriplate (MGB096-1-2-LG-L).

SxG absorbance and fluorescence spectra in the presence of λ-DNA were collected on a Tecan INFINITE M1000 PRO in a 96-Well Glass Bottom Matriplate (MGB096-1-2-LG-L). Absorbance spectra of SxG in the absence of DNA were collected on a Thermo Scientific Evolution 260 Bio UV-Visible Spectrophotometer in plastic PS cuvettes with a pathlength of 10 mm (BRAND Macro-Cuvettes, cat# 759030, Sigma-Aldrich).

We found a molar absorption coefficient of *ε*_529nm_ ≈ 90,000 M^-1^×cm^-1^ for SxO (Supplementary Figure S13ab) and of *ε*_500nm_ ≈ 74,000 M^-1^×cm^-1^ for SxG (Supplementary Figure S14ab). Absorbance and fluorescence spectra of SxO and SxG in DMSO (Dimethyl sulfoxide for molecular biology, cat# D8418, Sigma Aldrich) were collected in quartz cuvettes (QS, Hellma).

### Fluorescence lifetime measurements

We recorded time-correlated single-photon counting (TCSPC) data on a custom build instrument using a pulsed diode laser with excitation at 515 nm (Becker & Hickl BDL-515-SMN; controller: Becker & Hickl LSB-C laser switch box, 20 MHz laser repetition rate). Data are filtered through a long pass filter before signal collection. Finally, a microchannel plate photomultiplier tube (R3809U-50, 3.2kV, Hamamatsu) was used to detect single photons. Samples were prepared from identical SxO and DNA stocks as for the absorbance, excitation, and emission measurements and measured in the same 10 mm path length cuvettes. We measured the instrument response function (IRF) using the fast decay of Allura Red in an identical cuvette. The time resolution was determined as the FWHM of the IRF, which was measured to be 0.3 ns. All TCSPC data were analyzed using lifetime fitting software SPCImage 6.2 (Becker & Hickl).

### McGhee-von Hippel binding model

To describe the binding equilibrium of the intercalating dyes to DNA we used the cooperative McGhee-von Hippel binding isotherm ^35, 44, 45, 47^:

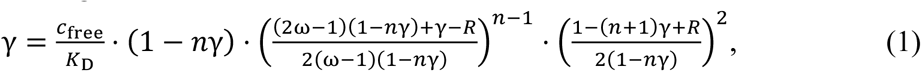

with,

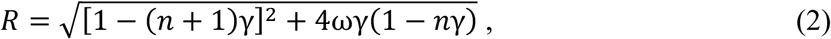

here, γ represents the binding density, i.e. the ratio of bound ligands to total base pairs, *c*_free_ is the free ligand concentration, *K*_D_ is the dissociation constant, *n* is the binding site, *ω* is a co-operativity parameter expressing the preference of a ligand binding to a singly or doubly contiguous site relative to an isolated site.

The McGhee-von Hippel binding model was fitted to our experimental data according to

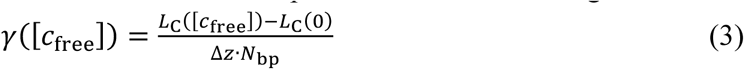

with *L*_C_([*c*_free_]) the DNA contour length, Δ*z* the increase in DNA contour length per ligand bound, and *N*_bp_ is the number of base pairs in the DNA construct (*N*_bp_ =21,000). We assumed a standard value for the DNA elongation per binding event, Δ*z* = 0.34 nm, which is common for intercalators^28, 35, 39, 47^.

Since the ends of the DNA are torsionally constrained the linking number *Lk* is constant for a given number of applied turns in the MT. By White’s formula (also known as Calugareanu-White-Fuller theorem ^26, 73, 74, 75^) *Lk = Tw + Wr,* i.e. the linking number *Lk* is the sum of twist *Tw* and writhe *Wr*. For constant *Lk*, a decrease in the *Tw* of DNA upon intercalation must be compensated by an increase in *Wr*, leading to the formation of positive plectonemes (and a corresponding decrease in tether extensions) if the dye concentration is increased while the number of applied turns is held constant. Consequently, after increase of SxO or SxG concentration, negative turns must be applied to return to a DNA conformation without plectonemes, i.e. maximum extension ^35, 39^. We quantify the twist change by determining the centers of the DNA rotation curves (Figure 3) and we describe the DNA twist change upon intercalation according to

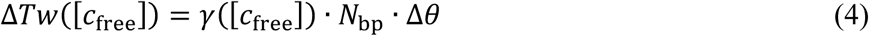

with Δ*Tw* the total change in DNA twist and Δ*θ* the unwinding angle per intercalation event. In our single-molecule magnetic tweezers assays, the DNA concentration is much lower than the dye concentration, [DNA]≪[SxO]. Therefore, the free ligand concentration is approximately equal to the total dye concentration *c*_total_, i.e. *c*_free_ ≈ [SxO] or *c*_free_ ≈ [SxG].

When the free ligand concentration is on the order of the bound ligand concentration, the assumption *c*_free_ ≈ *c*_total_ no longer holds ^39, 76^. In this case, the finite dye and DNA concentrations need to be included. Therefore, to apply the model to bulk optical spectroscopy methods, we implemented the McGhee-von Hippel model (Equation 1) by expressing the total ligand concentration as *c*_total_ = *c*_free_+ *C_bound_* ;and the binding density *γ* as *γ* = *c*_bound_/*c*_DNA_, where *c*_DNA_ is the total DNA concentration, as done in previous work.

To apply the model to the optical properties of the dye in the presence of varying DNA concentrations, we fitted *c*_bound_ to the data or computed *c*_bound_ using the binding parameters found through single-molecule MT manipulation. To compare the binding behaviors observed using MT and absorbance spectroscopy, we fitted a linear superposition of the free *A*_free_ and bound *A*_bound_ dye contributions to the absorbance intensity data:

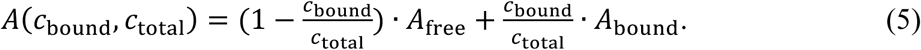

Analogously, to compare the model to the spectral shift upon binding, we fitted an empirical linear superposition of the wavelengths of the absorbance maxima of the free *λ*_max,free_ and bound *λ*_max,bound_ dye:

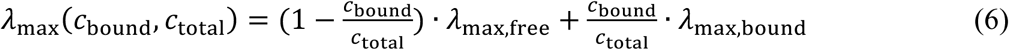

To adjust the model for the fluorescence data, we assume that the fluorescence intensity *I* increases linearly with the bound dye (since *I*≫*I*_0_) ^39^:

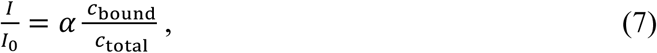

with *α* being an enhancement factor that accounts for the fluorescence enhancement of SxO upon binding to dsDNA. We neglect contributions of the free dye since the fluorescence intensities are much lower than in presence of DNA (see below). The fluorescence intensity data are compared to the MT data by combining Equations 1 and 7, while assuming *ω*=1 for low binding densities, and fitting to the fluorescence spectroscopy data.

### Quantum Chemical Calculations

Initial structures of the four molecules were built from scratch in TMoleX version 2026 ^77^. As a first step, the structures were optimized with density functional theory (DFT) calculations employing the B3LYP functional ^78, 79, 80^ and cc-pVTZ basis set ^81, 82^. Subsequent frequency calculations at the same level of theory exhibited only positive vibrational frequencies indicating that for each molecule a minimum on the potential energy surface was found. These DFT calculations were realized with Gaussian 16 ^83^. These structures were further optimized with the resolution-of-identity second-order approximate coupled-cluster (RI-CC2) method ^84, 85, 86^ employing cc-pVDZ as basis set together with the corresponding auxiliary basis set ^87^. Finally, RI-CC2 excited state calculations ^88^ for the first ten excited states for each molecule were performed employing the aug-cc-pVDZ basis set to improve the description of excited states. All RI-CC2 calculations were combined with the conductor-like screening model (COSMO) using the post-SCF reaction field scheme ^89^. To model water (DMSO) as solvent, a relative dielectric constant of 80.1 (46.7) and a refractive index of 1.33 (1.4783) was used. The RI-CC2 calculations were realized with Turbomole Version 8.0 ^90^. A tightened SCF convergence criterion of 10^-8^ and default frozen core orbitals were employed, otherwise default values were used. The molecular orbitals were visualized with Jmol.

## ACKNOWLEDGEMENTS

We thank Maria Kurilova, Sidonie Lieber, Andreas Spörhase, Olga Ustinov, and Elleke van Harten for laboratory support, Andries Meijerink, Patrick Ganswindt, and Alexander Urban for support with fluorescence measurements, and Pauline Kolbeck for useful discussions. We thank Caroline Körösy for preparation of the DNA construct for MT measurements and for laboratory assistance.

## AUTHOR CONTRIBUTIONS

Koen R. Storm: Conceptualization, Investigation, Software, Formal analysis, Visualization, Writing - original draft, Project administration. Stefanie D. Pritzl: Investigation, Software, Formal analysis, Project administration, Funding acquisition. Yi-Yun Lin: Investigation (MT), Formal analysis. Christian Wiebeler: Investigation (QCC), Formal analysis. Alptuğ Ulugöl: Software. Martin Lehmann: Investigation (MS). Dave van den Heuvel: Resources. Gerhard A. Blab: Formal analysis. Gerd Gemmecker: Investigation (NMR), Formal analysis (NMR). Jan Lipfert: Conceptualization, Software, Writing - original draft, Supervision, Funding acquisition. All authors contributed to review and editing of the final version.

## FUNDING

This work was supported by Utrecht University, by the European Research Council Consolidator Grant “ProForce” (101002656), the Horizon Europe 2021-2027 Framework Programme Grant Agreement number 101168851, and a Feodor-Lynen fellowship from the Alexander von Humboldt Foundation. The authors gratefully acknowledge the resources on the LiCCA HPC cluster of the University of Augsburg, co-funded by the German Research Foundation DFG – Project-ID 499211671 as well as financial support from the DFG via TRR-386, TP B06, project number 514664767. NMR measurements were acquired on the spectrometers of the Bavarian NMR Center (BNMRZ). Disclaimer: Funded by the European Union. Views and opinions expressed are however those of the author(s) only and do not necessarily reflect those of the European Union. The European Union cannot be held responsible for them.

## CONFLICT OF INTEREST

The authors declare no competing interests.

## SUPPLEMENTARY DATA

Supplementary data is available online.

## Notes

### Competing Interest Statement

The authors have declared no competing interest.

