## Supplementary Information for "Molecular Structure, DNA Binding, and Photophysical Properties of SYTOX Orange and SYTOX Green"

**Content:**
**Supplementary Tables S1-S3**
**Supplementary Figures S1-S21**
**Supplementary References**

### SUPPLEMENTARY TABLES

**Supplementary Table S1. Chemical shift table for SYTOX Orange and SYTOX Green.** The numbering scheme is given in the molecular structure at the end of the table. Chemical shifts for $^{1}H$ signals are given in normal font, for corresponding $^{13}C$ signals in italics; $^{15}N$ shifts are marked accordingly. For entries marked with a "*" the assignment is ambiguous.

| | SYTOX Green (DMSO, 298 K) | SYTOX Orange (DMSO, 298 K) |
|---|---|---|
| **<u>Quinoline</u>** | | |
| **q** | *138.9* | |
| **o** | *7.77* | |
| | 129.9 | |
| **m** | *7.84* | |
| | 131.3 | |
| **p** | *7.74* | |
| | 130.7 | |
| **N1** | 157.2 | 142.9 |
| **N1-CH$_3$** | | 3.89 |
| | | *40.0* |
| **2** | *157.9* | *151.0* |
| **3** | 7.13 | 8.43 |
| | *103.4* | *112.4* |
| **4** | *150.0* | *148.8* |
| **4a** | *122.5* | *124.90* |
| **5** | 8.77 | 7.87 |
| | *125.7* | *105.4* |
| **6** | 7.67 | |
| | *126.4* | *157.9* |
| **6-OCH3** | | 4.03 |
| | | *56.7* |
| **7** | 7.76 | 7.64 |
| | *133.4* | *123.8* |
| **8** | 7.11 | 8.09 |
| | *119.3* | *121.0* |
| **8a** | *133.4* | *134.8* |
| **<u>Benzo-X-azole</u>** | | |
| **2a′** | 6.95 | 6.54 |
| | *88.2* | *81.3* |
| **2′** | *160.1* | *171.9* |
| **N3'** | *143.3* | 204.5 |
| **3′-CH$_3$** | 4.06 | |
| | *34.4* | |
| **3a′** | 141.2 | *154.9** |

| | | | |
|---|---|---|---|
| **4’** | 7.82 | | |
| | *113.4* | **N4’** | 163.5 |
| | | **4’-$CH_3$** | 4.07 |
| | | | *40.2* |
| **5′** | 7.66 | | 8.21 |
| | *128.8* | | *135.9* |
| **6′** | 7.47 | | 7.30 |
| | *124.90* | | *116.2* |
| **7'** | 8.16 | | 8.07 |
| | *123.5* | | *118.1* |
| **7a'** | *124.0* | | *145.7** |

| | | SYTOX Green (DMSO, 298 K) | SYTOX Orange (DMSO, 298 K) |
|---|---|---|---|
| **<u>Tail (R')</u>** | | | |
| **α** | | | |
| | **Nα** | *72.0* | *137.5* |
| **β / β'** | | 3.26 | 7.85 |
| | | *49.6* | *130.0* |
| **γ / γ'** | | 1.74 | 7.83 |
| | | *21.0* | *134.0* |
| **δ / δ'** | | 3.19 | |
| | | *63.3* | *130.3* |
| **Nδ-$CH_3$** | | | |
| **ε** | | | 4.64 |
| | **Nε** | 49.2 | *63.5* |
| **Nε-$CH_3$** | | 3.03 | |
| | | *53.0* | |
| **Nζ** | | | 61.5 |
| **Nζ-$CH_3$** | | | 2.96 |
| | | | *46.7* |
| **η** | | | 3.31/3.41 |
| | | | *55.8* |
| **θ** | | | 1.37 |
| | | | *8.2* |

SYTOX Green

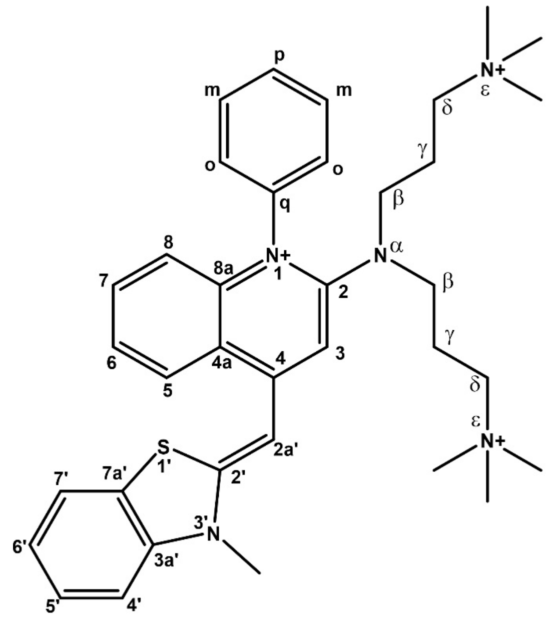

SYTOX Orange

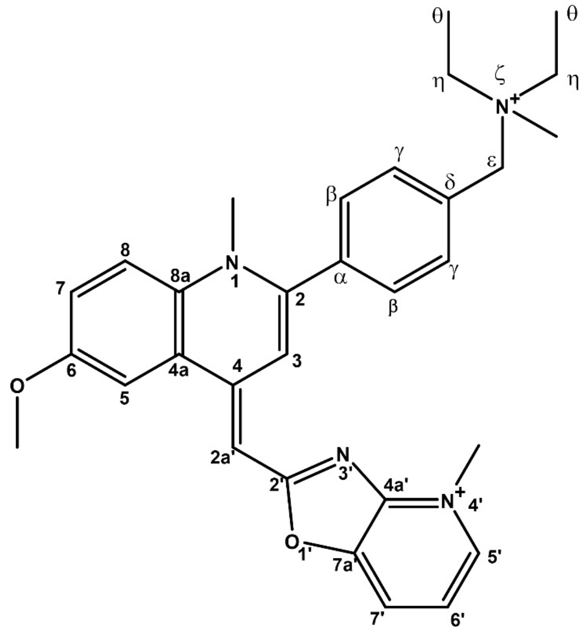

**Supplementary Table S2. Excitation energies (ΔE), wavelengths (λ), and oscillator strengths (f) for SxO, SxG, SYBR Gold, and PicoGreen in water.** Data for all four molecules were obtained from RI-CC2/aug-cc-pVDZ calculations employing the COSMO solvation model with parameters representing water as solvent.

| ΔE (eV) | λ (nm) | f |
|---|---|---|
| **SxO** | | |
| 2.39 | 519.5 | 1.110 |
| 3.52 | 352.5 | 0.025 |
| 3.65 | 339.9 | 0.059 |
| 3.95 | 314.0 | 0.037 |
| 4.25 | 291.4 | 0.365 |
| 4.39 | 282.5 | 0.014 |
| 4.45 | 278.5 | 0.069 |
| 4.61 | 268.8 | 0.222 |
| 4.72 | 262.5 | 0.093 |
| 4.80 | 258.3 | 0.015 |
| **SxG** | | |
| 2.49 | 497.5 | 0.909 |
| 3.37 | 368.2 | 0.233 |
| 3.96 | 312.9 | 0.003 |
| 4.15 | 299.0 | 0.044 |
| 4.20 | 294.9 | 0.090 |
| 4.32 | 286.8 | 0.133 |
| 4.36 | 284.6 | 0.131 |
| 4.45 | 278.5 | 0.003 |
| 4.67 | 265.3 | 0.021 |
| 4.69 | 264.2 | 0.039 |

| ΔE (eV) | λ (nm) | f |
|---|---|---|
| **SYBR Gold** | | |
| 2.55 | 485.5 | 0.983 |
| 3.60 | 344.7 | 0.077 |
| 3.92 | 316.5 | 0.024 |
| 4.27 | 290.3 | 0.184 |
| 4.39 | 282.5 | 0.369 |
| 4.48 | 276.5 | 0.034 |
| 4.65 | 266.6 | 0.070 |
| 4.69 | 264.4 | 0.089 |
| 4.75 | 260.8 | 0.150 |
| 4.83 | 256.9 | 0.082 |
| **PicoGreen** | | |
| 2.52 | 492.5 | 0.969 |
| 3.41 | 363.4 | 0.230 |
| 4.02 | 308.8 | 0.002 |
| 4.17 | 297.4 | 0.041 |
| 4.22 | 293.5 | 0.115 |
| 4.34 | 285.5 | 0.076 |
| 4.36 | 284.2 | 0.162 |
| 4.47 | 277.6 | 0.005 |
| 4.56 | 271.7 | 0.001 |
| 4.72 | 262.8 | 0.018 |

**Supplementary Table S3. Excitation energies (ΔE), wavelengths (λ), and oscillator strengths (f) for SxO, SxG, SYBR Gold, and PicoGreen in DMSO.** Data for all four molecules were obtained from RI-CC2/aug-cc-pVDZ calculations employing the COSMO solvation model with parameters representing DMSO as solvent.

| ΔE (eV) | λ (nm) | f |
|---|---|---|
| **SxO** | | |
| 2.34 | 530.4 | 1.138 |
| 3.48 | 356.0 | 0.024 |
| 3.64 | 340.9 | 0.064 |
| 3.94 | 314.6 | 0.043 |
| 4.24 | 292.6 | 0.409 |
| 4.39 | 282.4 | 0.013 |
| 4.44 | 279.1 | 0.068 |
| 4.60 | 269.6 | 0.246 |
| 4.72 | 262.8 | 0.096 |
| 4.80 | 258.5 | 0.019 |
| **SxG** | | |
| 2.45 | 505.6 | 0.938 |
| 3.36 | 369.2 | 0.251 |
| 3.96 | 313.3 | 0.003 |
| 4.14 | 299.5 | 0.071 |
| 4.19 | 296.0 | 0.086 |
| 4.32 | 287.2 | 0.136 |
| 4.34 | 285.4 | 0.138 |
| 4.45 | 278.4 | 0.003 |
| 4.67 | 265.6 | 0.025 |
| 4.69 | 264.3 | 0.043 |

| ΔE (eV) | λ (nm) | f |
|---|---|---|
| **SYBR Gold** | | |
| 2.51 | 494.6 | 1.008 |
| 3.58 | 346.3 | 0.080 |
| 3.91 | 317.4 | 0.024 |
| 4.26 | 291.1 | 0.223 |
| 4.37 | 283.4 | 0.395 |
| 4.48 | 276.5 | 0.033 |
| 4.64 | 267.0 | 0.076 |
| 4.68 | 264.8 | 0.109 |
| 4.75 | 261.1 | 0.151 |
| 4.81 | 257.6 | 0.073 |
| **Pico Green** | | |
| 2.48 | 500.6 | 0.997 |
| 3.40 | 364.4 | 0.249 |
| 4.01 | 309.2 | 0.002 |
| 4.16 | 297.9 | 0.067 |
| 4.21 | 294.6 | 0.114 |
| 4.34 | 285.9 | 0.059 |
| 4.35 | 285.0 | 0.187 |
| 4.47 | 277.4 | 0.005 |
| 4.55 | 272.4 | 0.001 |
| 4.72 | 262.9 | 0.020 |

### SUPPLEMENTARY FIGURES

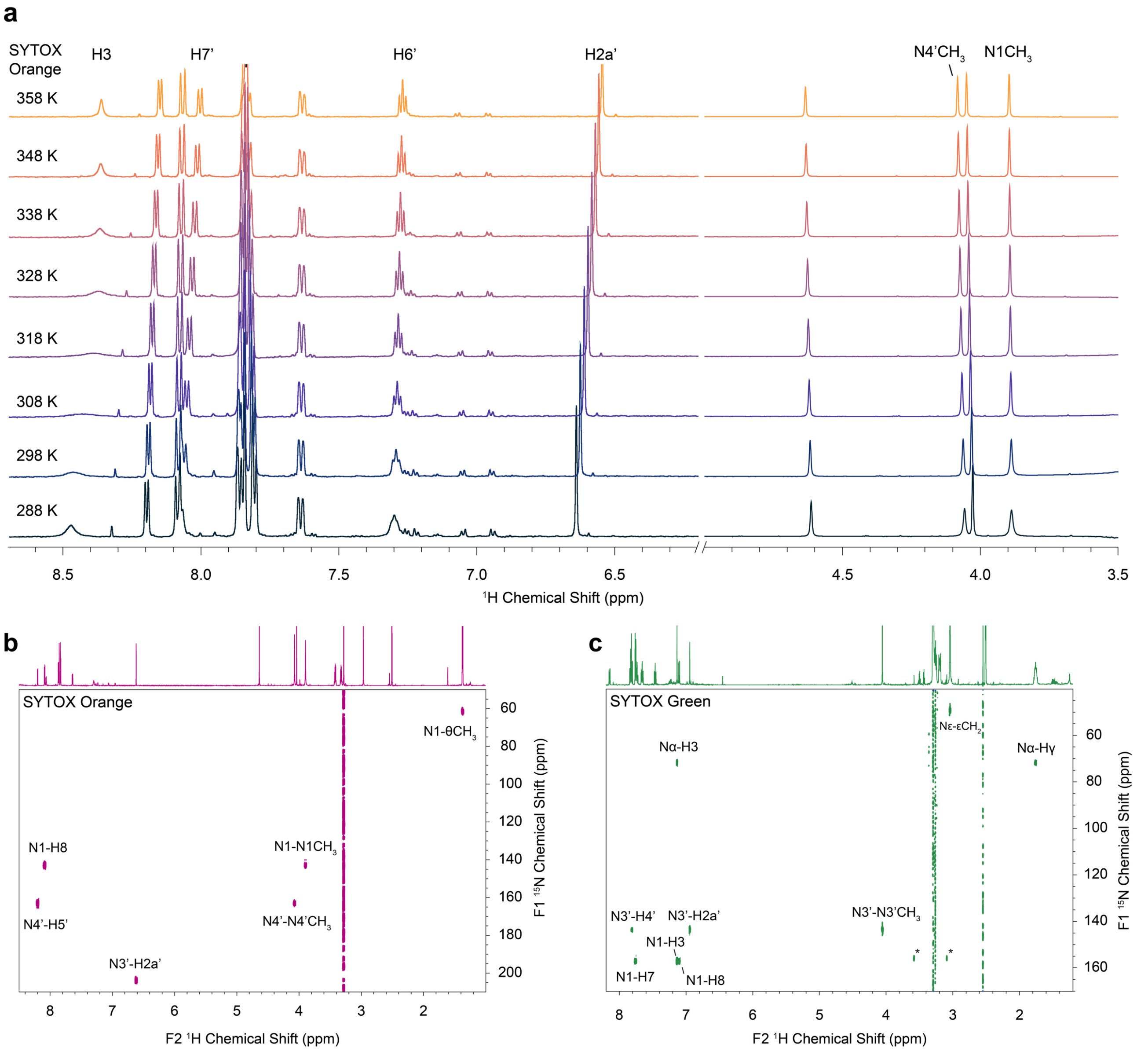


**Supplementary Figure S1. NMR-based structure elucidation of SxO and SxG.** a) $^{1}H$ NMR spectra of SYTOX Orange as a function of temperature in the range 288−358 K in DMSO-d$_6$. b) $^{1}H,^{15}N$-HMBC spectrum of SYTOX Orange, showing the assignment of all four nitrogens from long-range correlations, including the unusual N3' shift of ca. 205 ppm. c) $^{1}H,^{15}N$-HMBC spectrum of SYTOX Green with assignment of the four different nitrogen signals from long-range correlations.

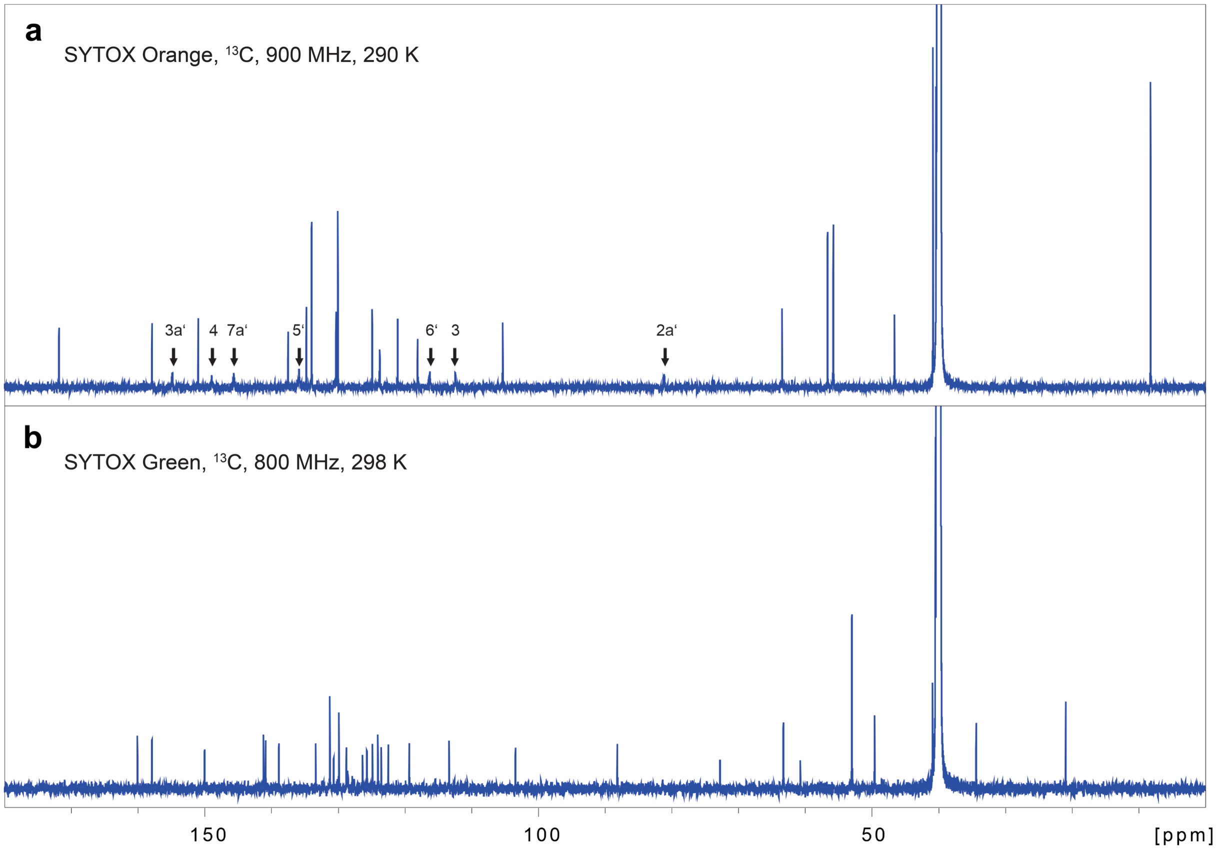


**Supplementary Figure S2. 1D $^{13}$C NMR spectra of SxO and SxG at 298 K.** a) $^{13}$C NMR spectrum of SxO (800 MHz $^{1}$H frequency); reduced intensity signals broadened from dynamic effects are marked and assigned (arrows). Inspection of the molecular structure of SxO shows that most of them belong to the aza-benzoxazol moiety, and they are all centered around the methine bridge. This strongly hints to a temperature-sensitive equilibrium between conformers caused by hindered rotation at this position. b) $^{13}$C NMR spectrum of SxG (measured at 900 MHz $^{1}$H frequency). Here signal intensities are uniform and no temperature-dependent line broadening was observed.

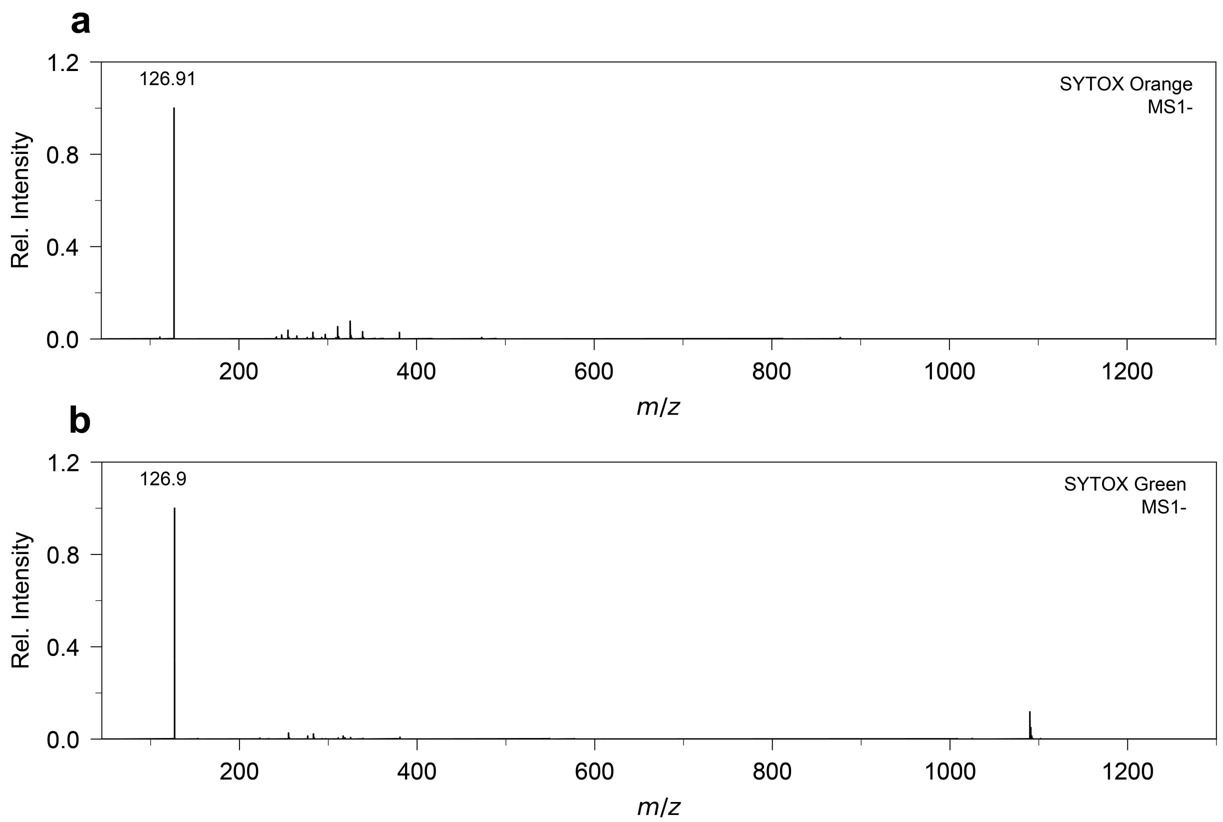


**Supplementary Figure S3. ESI-MS spectrum of SxO and SxG.** Mass spectra of a) SxO and b) SxG in negative mode indicating the presence of iodide as the relevant anion for both dyes.

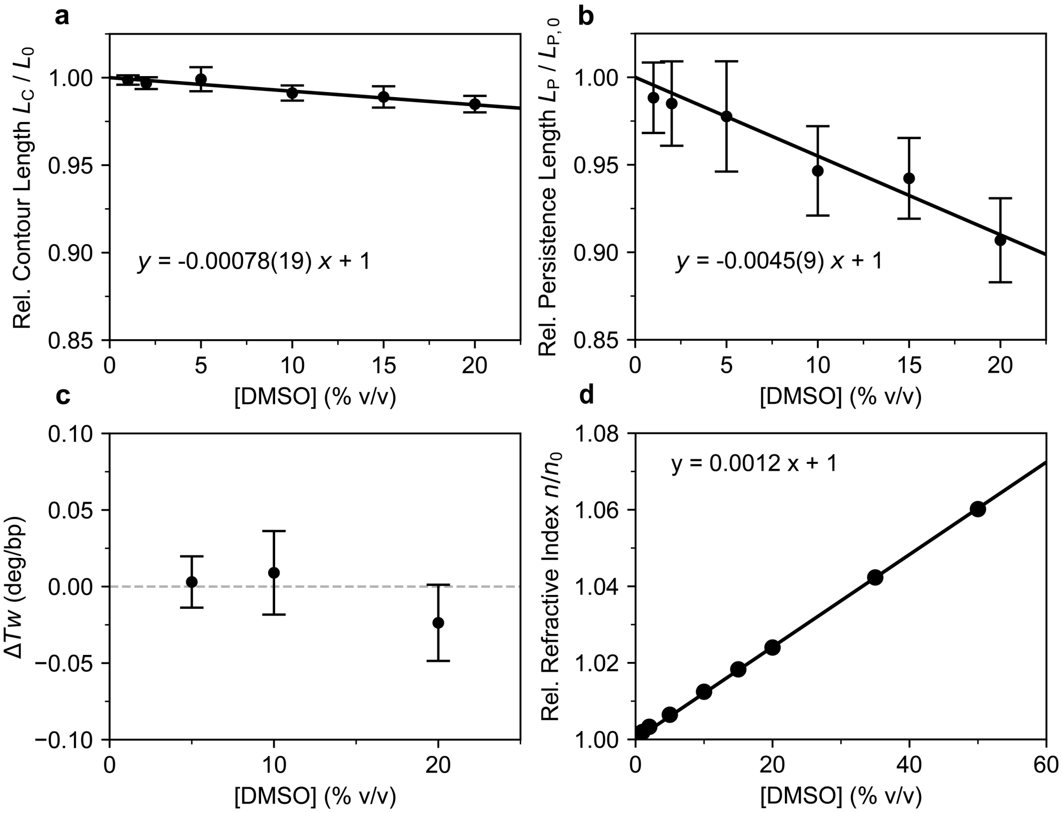


**Supplementary Figure S4. Effect of DMSO on MT measurements.** We test the effects of DMSO in the buffer, which is added upon adding SxO or SxG since the stocks are stored in DMSO, by performing MT measurements using the same set up and DNA tethers used for the dye measurements using PBS buffer with different concentrations of DMSO added. a) Contour length $L_C$ as a function of DMSO concentration obtained from fits of the WLC model to MT force-extension data. Data are normalized to the fitted contour length in the absence of DMSO (i.e. in PBS buffer). The solid line is a linear fit to the data ($R^2 = 0.99$). b) Bending persistence length $L_P$ as a function of DMSO concentration obtained from fits to the WLC model. Data are normalized to the fitted persistence length in the absence of DMSO. The solid line is a linear fit to the data ($R^2 = 0.98$). c) Change in the DNA twist as a function of DMSO concentration as determined from the centers of rotation curves. The twist is relative to DNA in the absence of DMSO. The data indicate essentially no change in twist upon addition of DMSO. d) Refractive index as a function of DMSO concentration in PBS buffer, determined using an Abbe refractometer (Abbe Refractometer 3T, Atago) at a wavelength of λ=589.3 nm and temperature of 21 ºC. Data are relative to the refractive index of water and in good agreement with the measurements by LeBel and Goring at 546 nm in water [1]. The solid line is a linear fit to the data ($R^2 = 0.999$). The data suggest that the direct effects of DMSO alone on the length, bending stiffness, and in particular twist of DNA are essentially negligeable in the range introduce by adding SxO or SxG dyes (< 20%, typically < 1%), fully consistent with a recent detailed investigation of DNA on the presence of DMSO [2]. Nonetheless, DMSO can have second order effects via altering the affinity of the dyes to DNA, see below.

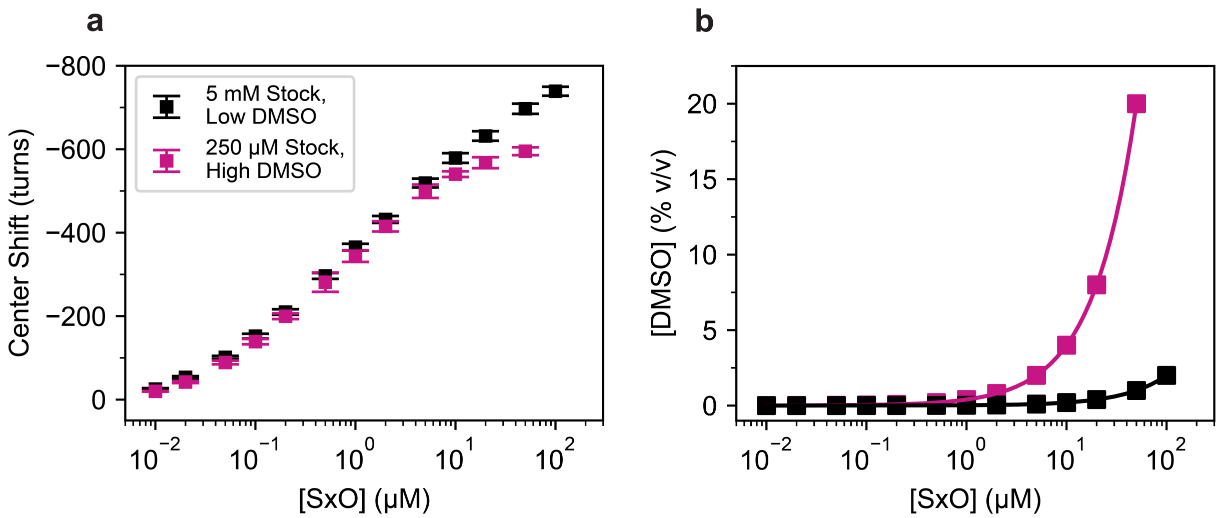


**Supplementary Figure S5. Effect of DMSO on SxO binding behavior revealed by MT rotation-extension measurements.** a) DNA unwinding quantified from the shift in the center position of the rotation extension curves for different SxO concentrations, using two different stocks that result in a low or a high DMSO background. SxO concentrations with a low DMSO background were prepared using a concentrated stock solution: 5 mM SxO in DMSO (Invitrogen, S11368) (black symbols). SxO concentrations with a high DMSO background were prepared using a less concentrated stock solution: 250 µM SxO in DMSO (Invitrogen, S34861) (red symbols). b) Percentage of DMSO present in the SxO supplemented buffer solutions. Lines and symbols are computed for SxO dilutions from the 5 mM or 250 µM SxO stock (same color code as panel A). Data in panel a are mean ± SD from at least 8 independent molecules. Since the direct effect of DMSO on DNA twist is negligeable in the range of DMSO concentrations used (see above and Ref. [2]), we attribute the observed differences to changes to the DNA affinity and binding pattern of SxO in the presence of high concentrations of DMSO. To avoid complications due to the presence of DMSO, all measurements reported in the main text employed the high dye concentration stocks, thus limiting the DMSO concentration to ≤ 2%.

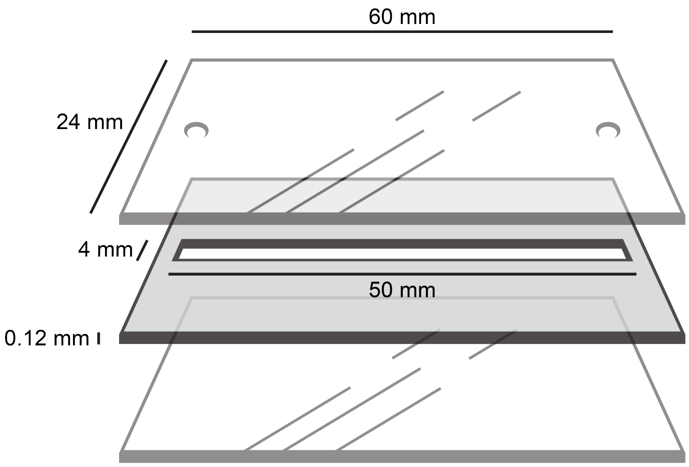


**Supplementary Figure S6. MT flow cell design.** Flow cells are prepared using two microscope coverslips (24x60 mm #1, Menzel). The bottom coverslip was functionalized with 3-Glycidoxypropyltrimethoxysilane and supplemented with polystyrene beads used as reference beads. To allow for liquid exchange, the top coverslip was prepared with two openings with a 1 mm radius. The flow cell is assembled by stacking the two coverslips using a layer of Parafilm (0.12 mm) with a central 4 x 50 mm channel cut out, resulting in a channel volume of ~ 25 μL. This channel is connected to the two openings of the top coverslip functioning as an inlet and an outlet for fluid exchange.

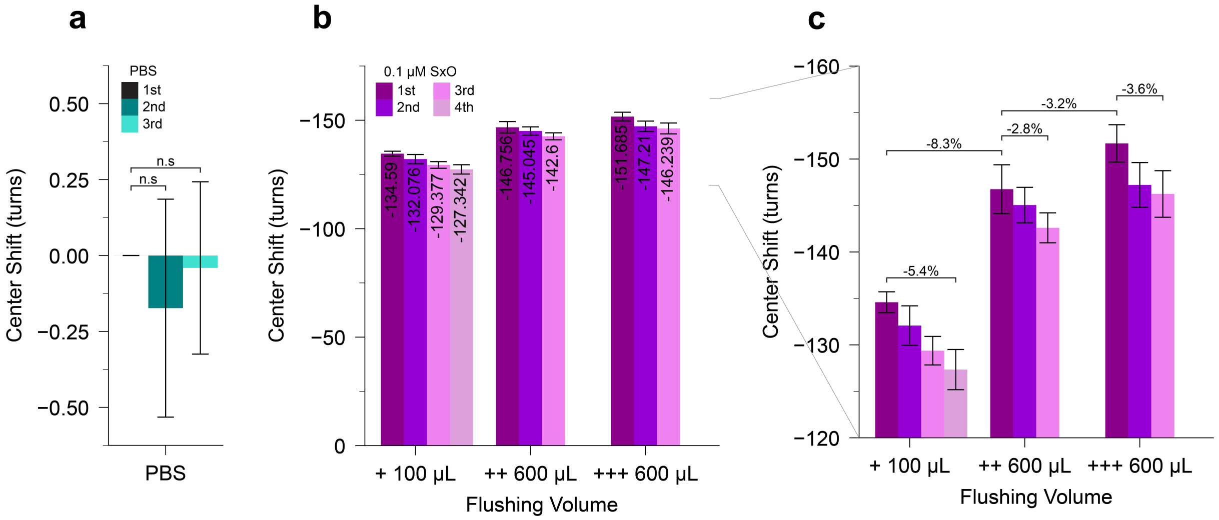


**Supplementary Figure S7. Equilibration of SxO concentration and binding in MT rotation-extension measurements.** To test the equilibration of dye concentration and repeatability of MT measurements, we performed repeated rotation-extension curve measurements under various conditions and determined the centers of the curves similar to the data shown in Figure 3. a) Rotation-extension curve centers for repeat measurements in PBS buffer. No significant difference is observed between repeats and curves are found to shift by less than one turn. b) and c) Shifts in the center position of rotation-extension curves after flushing different volumes of 0.1 µM SxO in PBS solutions. Values are relative to the center position of the rotation curve in PBS in the absence of SxO. First, rotation-extension measurements were performed in PBS. Subsequently, the flow cell (Supplementary Figure S6) was flushed with 100 µL (~4 cell volumes) of 0.1 µM SxO and four rotation extension curves were measured in series (taking ~8 min each). Afterwards, an additional 600 µL (~24 cell volumes) of 0.1 µM SxO was flushed through the flow cell (during a period of ~ 15 min) and three rotation extension curves were measured (taking again ~ 8 min each). This was repeated with an additional 600 µL of 0.1 µM SxO. Panel c is a close up of the data in panel b. Data in all panels are the mean ± std obtained from at least 7 DNA molecules.

The observed unwinding angle increases, albeit slightly, after subsequent flushed and decreases, again slightly, in repeated measurements after introducing a given volume of SxO. Small increases in apparent binding after subsequent flushes are likely due to somewhat incomplete fluid exchange in the flow cell, due to a surface boundary layer or a small dead volume. Small decreases in the apparent binding in subsequent measurements after flushing likely stem from some amount of surface sticking or absorption of dyes, similar what has been observed in previous measurements with intercalators [3, 4]. We note that the data imply that the kinetics of binding to DNA is considerably faster than the ~min time scale of the rotation curve measurements, as otherwise subsequent measurements after flushing would result in slow increases and not decreases in unwinding angle, consistent with previous kinetic measurements [5]. All data measurements reported in the main text flushed at least 600 µl solution at a given dye concentration, to keep the error due to incomplete equilibration to < 5%.

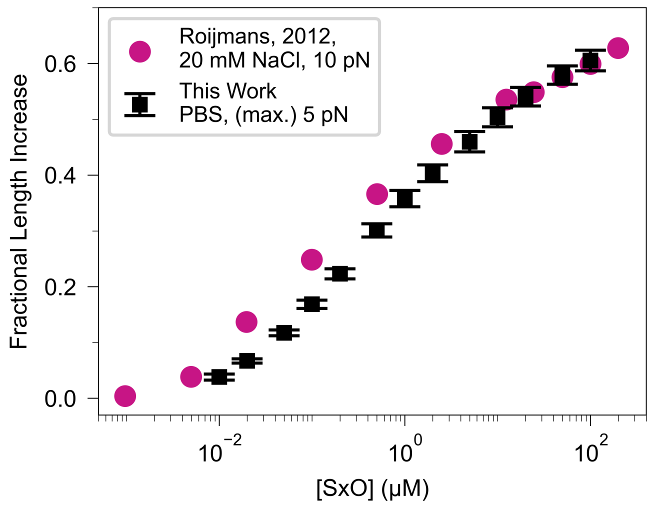


**Supplementary Figure S8. Comparison of SxO induced DNA lengthening measured using magnetic tweezers and optical tweezers.** Fractional length increase as function of SxO concentration obtained from the relative change in DNA contour length from fits of the WLC to MT force extension data ($F_{max}$=5 pN) (black squares) (same data as Fig 1). Fractional length increase as function of SxO concentration obtained measured extension at $F$=10 pN using optical tweezers (Data from Roijmans, 2012). The difference in the binding midpoint for different buffer conditions is likely explained by the salt dependence of the $k_{on}$ and $k_{off}$ rates [6] (20 mM monovalent salt for the Roijmans data; 140 mM monovalent salt for our measurements). DNA lengthening in the SxO concentrations range approaching saturation are within error. The data from Roijmans, 2012, are taken from "Characterization of the double stranded DNA stain SYTOX orange", Roijmans, R.F.H., 2012, Master thesis, TU Eindhoven, accessed from https://pure.tue.nl/ws/files/46932625/759029-1.pdf.

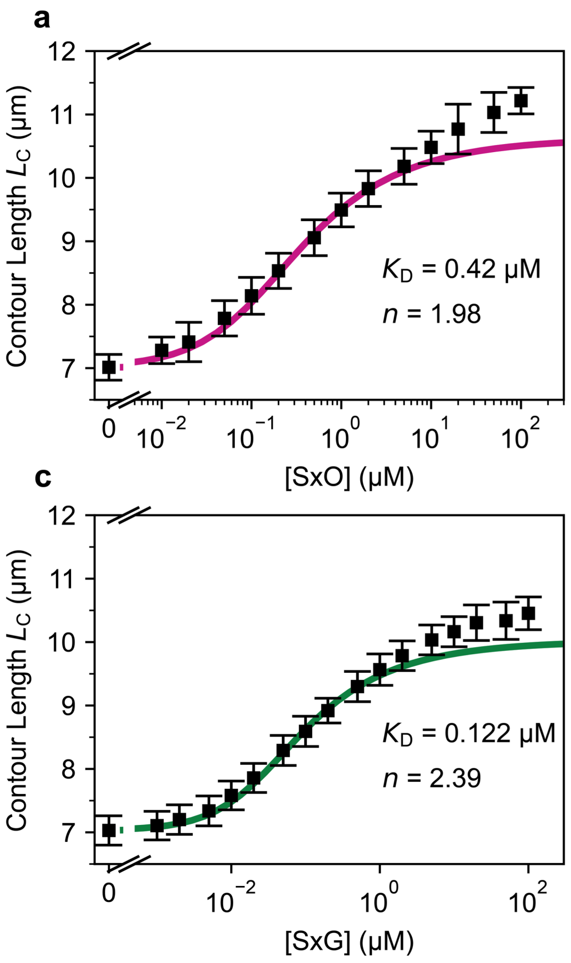


**Supplementary Figure S9. Fits of the (non-cooperative) McGhee-Von Hippel model to the concentration data.** a) DNA contour length as a function of SxO concentration (same data points as shown in Figure 2d) and fit of the non-cooperative McGhee-von Hippel model (solid line) to the data up to 2 µM SxO. b) Shifts of the centers of the rotation-extension curves as a function of SxO concentration (same data points as Figure 3b) and fit of the non-cooperative McGhee-von Hippel model (solid line), again up to 2 µM SxO.We find a dissociation constant $K_D$ = 0.42 µM and a binding site size $n$ = 1.98 for SxO. c) DNA contour length as a function of SxG concentration (same data points as shown in Figure 2g) and fit of the non-cooperative McGhee-von Hippel model (solid line) to the data up to 0.5 µM SxG. d) Shifts of the centers of the rotation-extension curves as a function of SxG concentration (same data points as Figure 3e), again up to 0.5 µM SxG, and fit of the non-cooperative McGhee-von Hippel model (solid line). For SxG, we find $K_D$ = 0.122 ± µM and $n$ = 2.39.

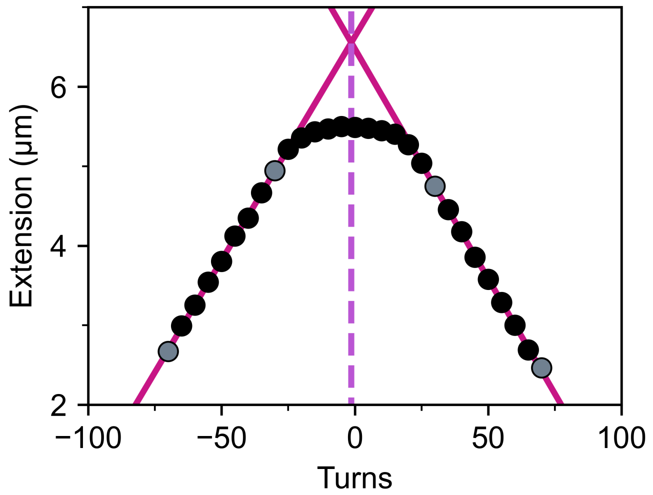


**Supplementary Figure S10. Fitting procedure for the MT rotation-extension curves.** The center shifts and slopes in the plectonemic regime of the rotation curves are determined by fitting two lines (dark red) to data in the linear post-buckling plectonemic regimes. The slope of the fitted lines are reported in Figure 3c,f. The centers of the rotation curves are determined as the intersection point of the two lines (shown as a vertical dashed line here).

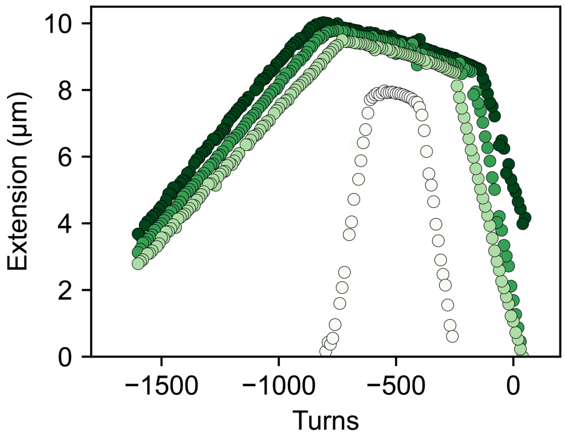


**Supplementary Figure S11 Stabilization of dsDNA by SxG for rotation-extension curves at elevated forces.** Rotation-extension curves for 21 kbp DNA in 20 µM SxG at different stretching forces. The exerted forces are (light to dark colors): 0.5, 3, 5 and 7 pN. Scattering is a measurement artifact and can be ignored. Occasional outlier points due to tracking error were removed for clarity.

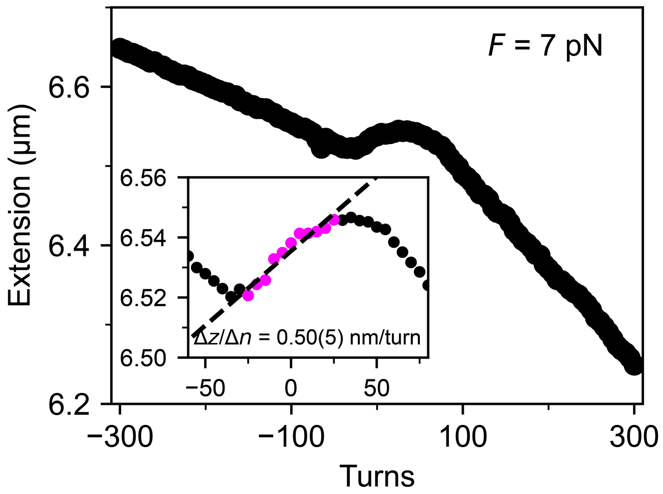


**Supplementary Figure S12 Rotation-extension curve of DNA at high force.** Detailed rotation-extension curve for DNA in PBS buffer at 7 pN. Note the difference in scale on the y-axis compared to the overview in Figure 3. The data show a region of increasing extension with increasing numbers of turns around zero turns, which is characteristic of the twist-stretch coupling of DNA [7, 8, 9]. Going to more positive or more negative turns, the extension decreases with increasing number of turns, due to melting [10] (at negative turns) and P-DNA formation [11] (at positive turns). The Inset shows a linear fit to the twist-stretch range indicated in violet. The value shown is the best fitting parameter of the slope and in brackets the error obtained from the covariance matrix of the fit and in excellent agreement with previous reports [7, 8, 9, 12].

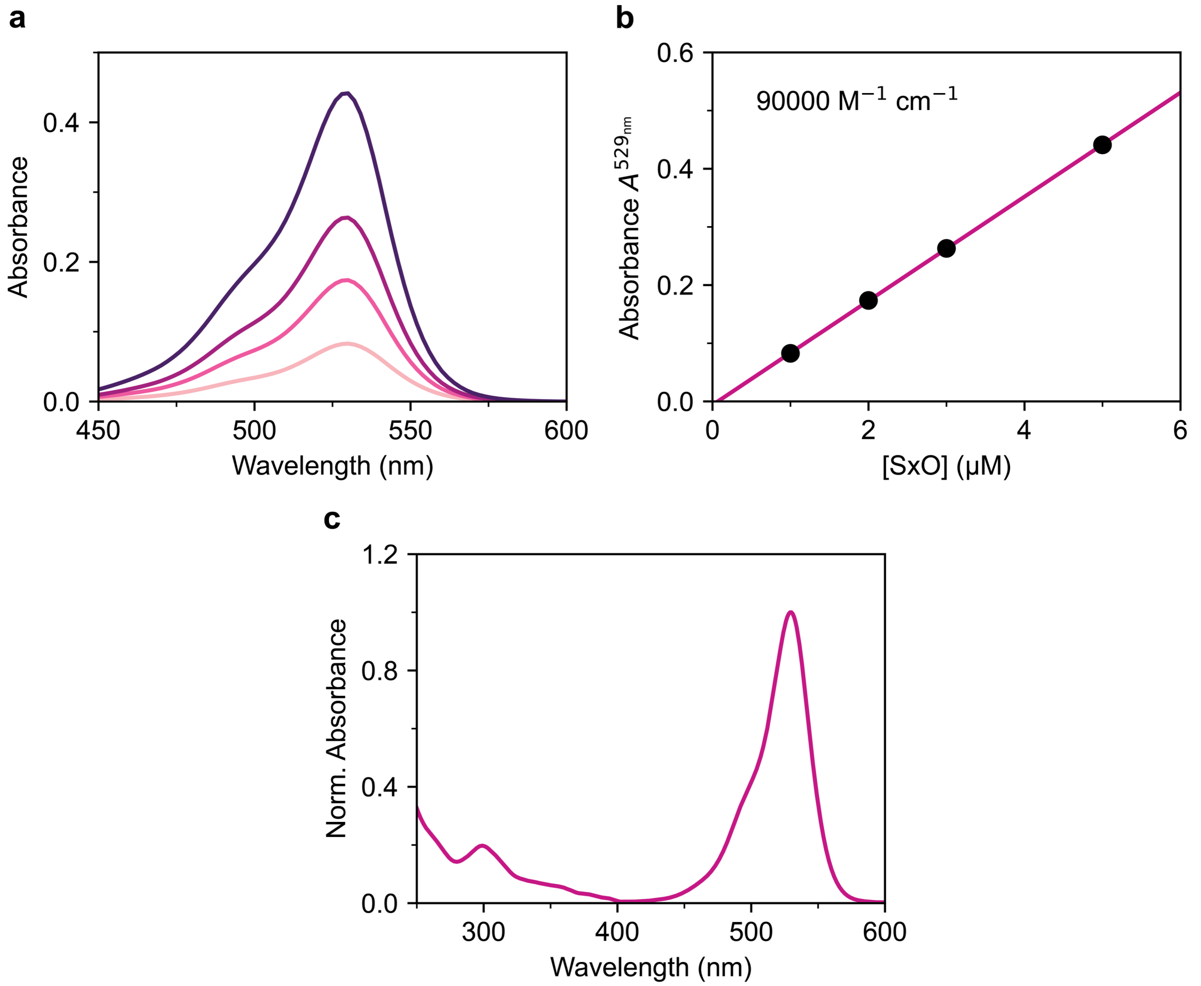


**Supplementary Figure S13. Absorbance measurements of SxO.** a) Absorbance spectra for a titration of SxO in PBS in the absence of DNA. The concentrations are (from light pink to purple) 1, 2, 3, 5 µM SxO. b) Absorbance value at a wavelength of $\lambda$=529 nm. The solid line is a linear fit according to the Beer-Lambert law, resulting in a molar absorption coefficient of $\varepsilon^{529nm} \approx$ 90,000 $M^{-1}$ $cm^{-1}$. c) Absorbance spectrum of 2.0 µM SxO in PBS in the visible and UV range. The main absorbance maximum at ~530 nm is clearly visible, with a minor peak at 300 nm.

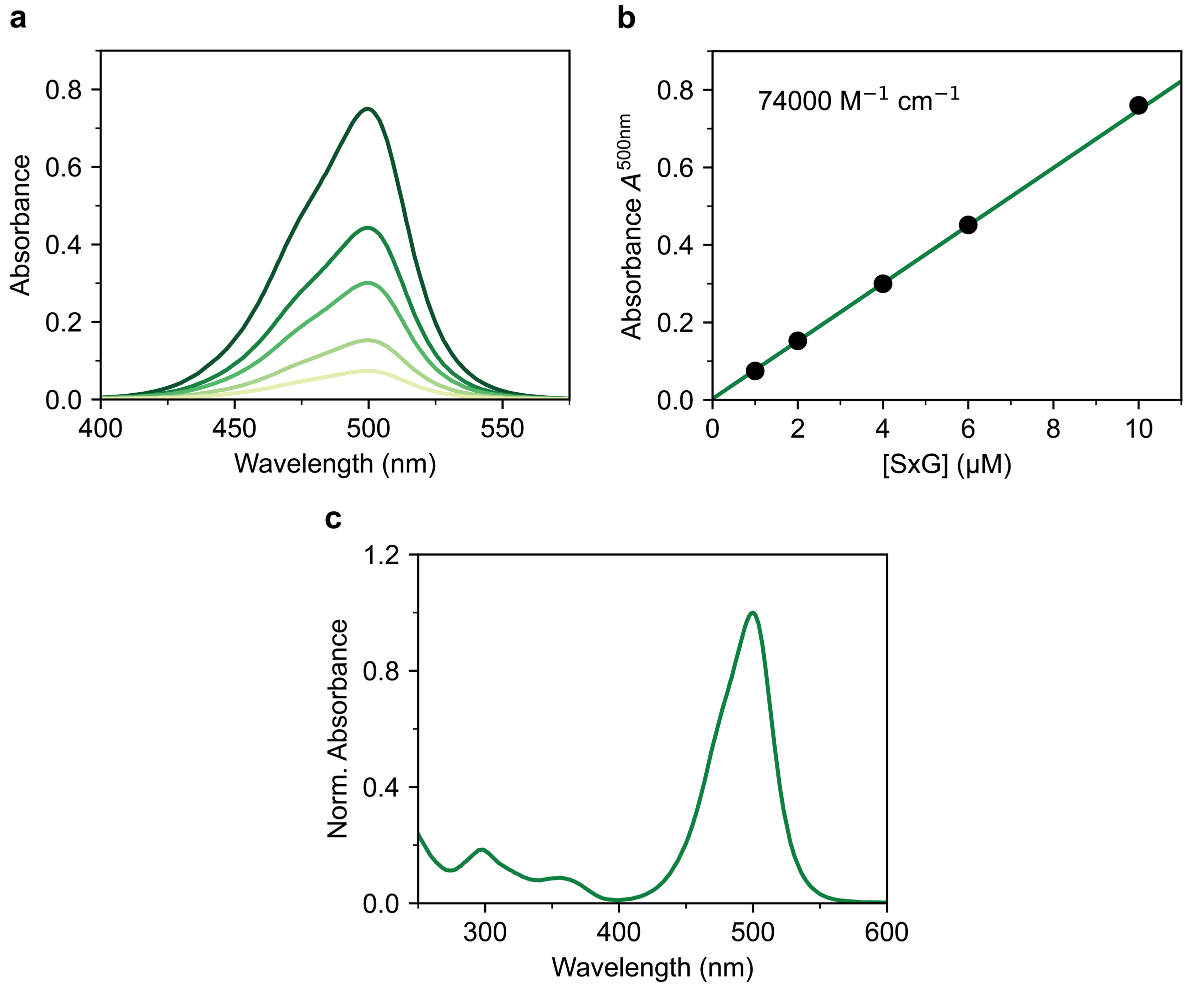


**Supplementary Figure S14. Absorbance measurements of SxG.** a) Absorbance spectra for a titration of SxG in PBS in the absence of DNA. The concentrations are (from light yellow to green,) 1, 2, 4, 6, 10 μM SxG. b) Absorbance values at a wavelength of $\lambda$=500 nm. The solid line is a linear fit according to the Beer-Lambert law, resulting in a molar absorption coefficient of $\varepsilon^{500nm} \approx$ 74,000 $M^{-1}$ $cm^{-1}$. c) Absorbance spectrum of 4 μM SxG in PBS in the visible and UV range. The main absorbance maximum at ~500 nm is clearly visible, with minor peaks around 300 and 360 nm.

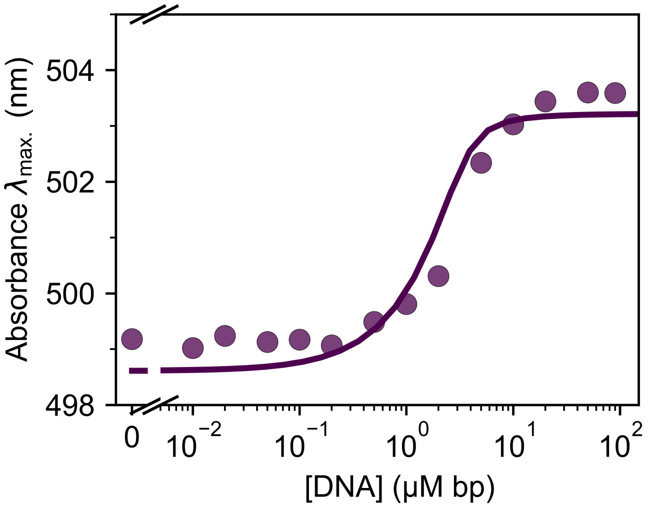


**Supplementary Figure S15. Alternative fit of the shift in the wavelength of the absorbance maximum for SxG upon binding to DNA.** Fit of the finite DNA concentration McGhee-von Hippel model to the wavelengths of maximum absorbance of SxG using the binding parameters $K_d$ and $n$ determined from the MT measurements. We find values of $\lambda_{max,free}$=498.6 nm and $\lambda_{max,bound}$=503.2 nm. Data points are the same as data in the inset of Figure 5d.

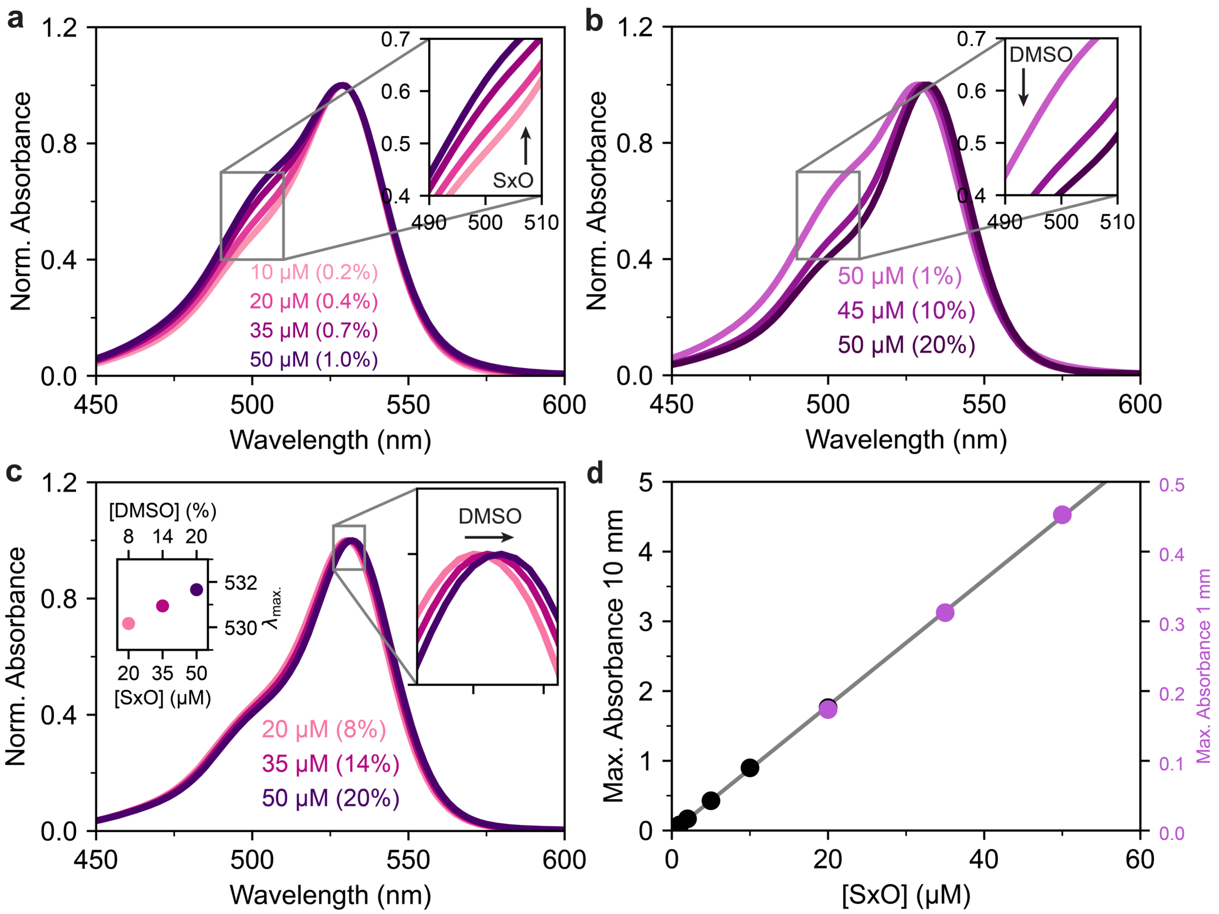


**Supplementary Figure S16. Absorbance spectra at high concentrations of SxO.** a) Absorbance spectra recorded at high concentrations of SxO (10-50 µM) and with low concentrations of DMSO (≤1% v/v) in PBS. The spectra are characterized by a single maximum at 528 nm and a shoulder around 500 nm. However, at high dye concentrations, the relative contribution of the band at ≈500 nm increases, which is suggestive of (self-)aggregation of SxO. Values in brackets indicate DMSO percentage. The spectrum of 50 µM SxO remained unchanged in the span of three days (data not shown). Pathlength was 0.1 cm, except for 10 µM where the pathlength was 1 cm. b) Absorbance spectra of 45-50 µM SxO with different amounts of DMSO. The contribution at ≈500 nm is reduced by addition of DMSO, suggesting a relationship between spectral deformation and solubility. Pathlength was 0.1 cm. c) Absorbance spectra of high concentrations of SxO (20-50 µM) with moderate concentrations of DMSO (≤20% v/v) in PBS. At high dye concentrations, no spectral deformation occurs. However, a slight 2 nm bathochromic shift of the entire spectrum is observed, most likely by the increasing concentration of DMSO. Pathlength was 0.1 cm. d) Maximum of the absorbance of SxO with moderate concentrations of DMSO (≤20% v/v) in PBS. The magnitude of maximum absorbance remains linear.

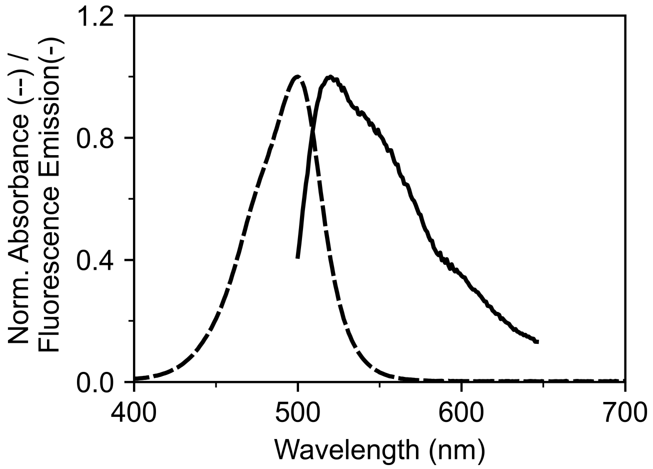


**Supplementary Figure S17. Absorbance and fluorescence emission spectra of SxG in PBS.** Concentration was 1 μM SxG in PBS. Absorbance was measured on a Thermo Scientific Evolution 260 Bio UV-Visible Spectrophotometer. Fluorescence emission was collected on a Horiba Fluoromax Plus Spectrofluorometer and background corrected.

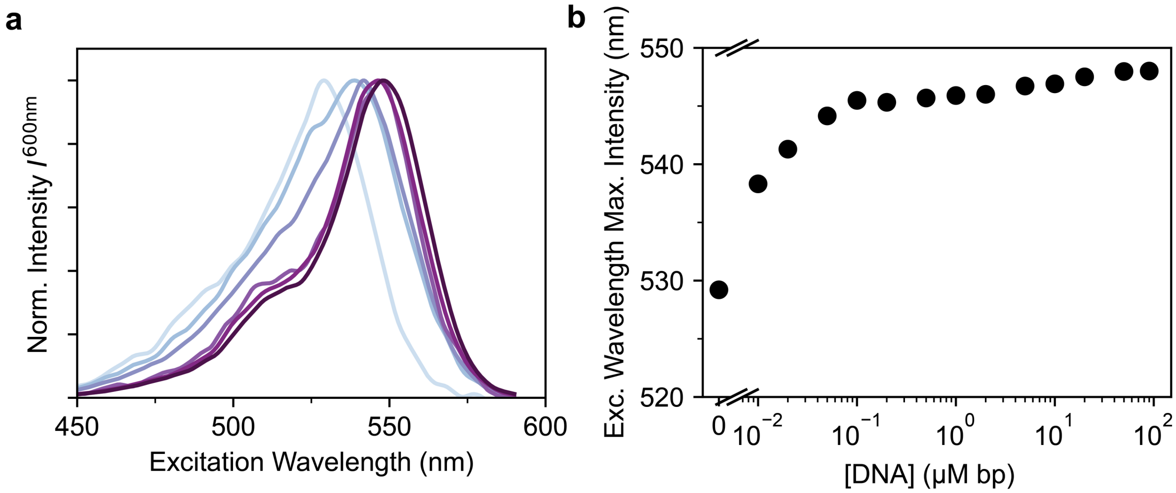


**Supplementary Figure S18. Fluorescence excitation spectra of SxO with increasing DNA concentration.** a) Fluorescence excitation spectra of SxO upon titration with λ-DNA. Measurements use a fixed concentration of 1.8 μM SxO in PBS solution. The base pair concentrations shown are (light to dark) 0, 0.01, 0.02, 0.1, 1, 90 μM. b) Excitation wavelength of maximum intensity. The emission intensity was collected at a wavelength of $\lambda_{em}$=600 nm.

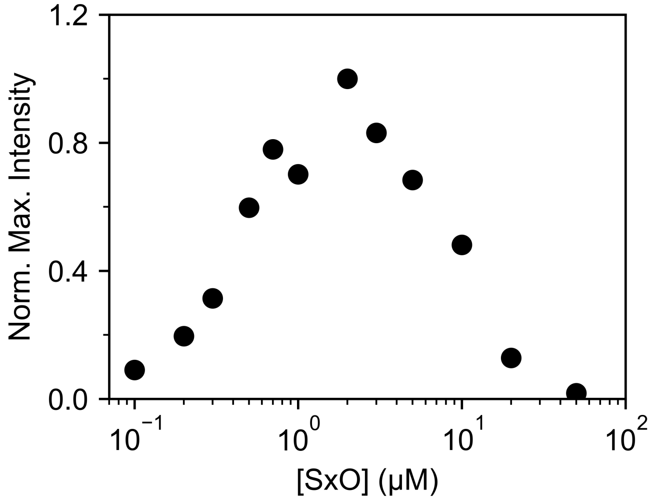


**Supplementary Figure S19. Fluorescence emission intensity upon increasing SxO concentration against a fixed DNA background.** Fluorescence emission intensity (after excitation at 530 nm) for a titration of SxO into a background of 2 μM·bp λ-DNA in PBS. The initial increase in intensity is due to the increase in DNA-bound SxO. However, the measured intensity plateaus and then decreases with increasing SxO concentrations above 2 μM, at least partially due to strong absorption and the inner filter effect. Data were obtained with a pathlength of 1 cm.

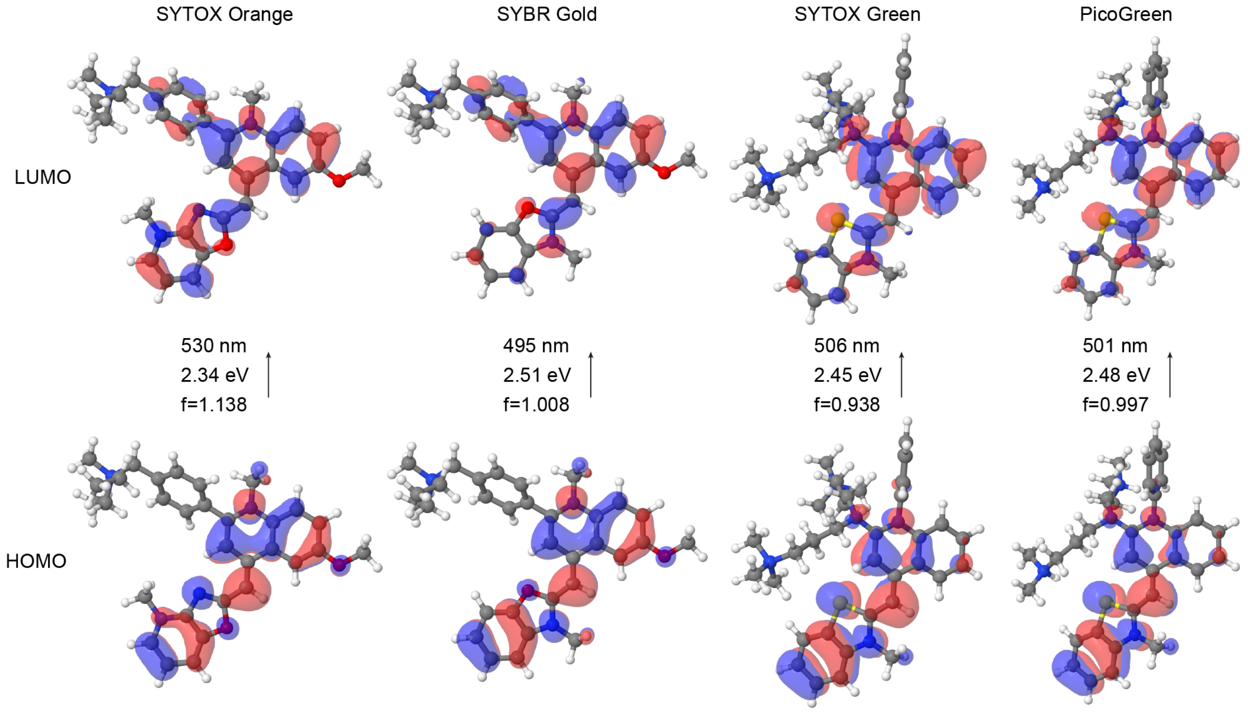


**Supplementary Figure S20. Quantum chemical calculations predict ground state structures and absorption wavelengths in DMSO.** Structures for SxO, SYBR Gold, SxG, and PicoGreen were optimized at the RI-CC2/cc-pVDZ level of theory and the subsequent excited state calculations with RI-CC2/aug-cc-pVDZ employing the COSMO solvation model with parameters representing DMSO as solvent were used for orbital visualization (Materials and Methods). The highest occupied molecular orbitals (HOMO; bottom) and lowest unoccupied molecular orbitals (LUMO; top) are shown. The excitation energies form the ground to the first excited state, the corresponding wavelengths, and the oscillator strengths are indicated for each dye. For each of the four molecules, this excitation is dominated by a transition from HOMO to LUMO. The orbitals and corresponding predicted absorption wavelengths are rather similar for SxG and PicoGreen, in agreement with experimental data and consistent with the very similar core structure. For SxO and SYBR Gold, the orbitals show clear differences, in particular for the LUMO, and the predicted absorption wavelength is considerably red shifted for SxO. Simulations using solvent parameter for water are shown in Figure 7. The predicted absorption wavelengths are shifted to longer wavelength in DMSO compared to aqueous solvent for all dyes investigated, in (semi-quantitative) agreement with our experimental findings for SxO and SxG (Table 2).

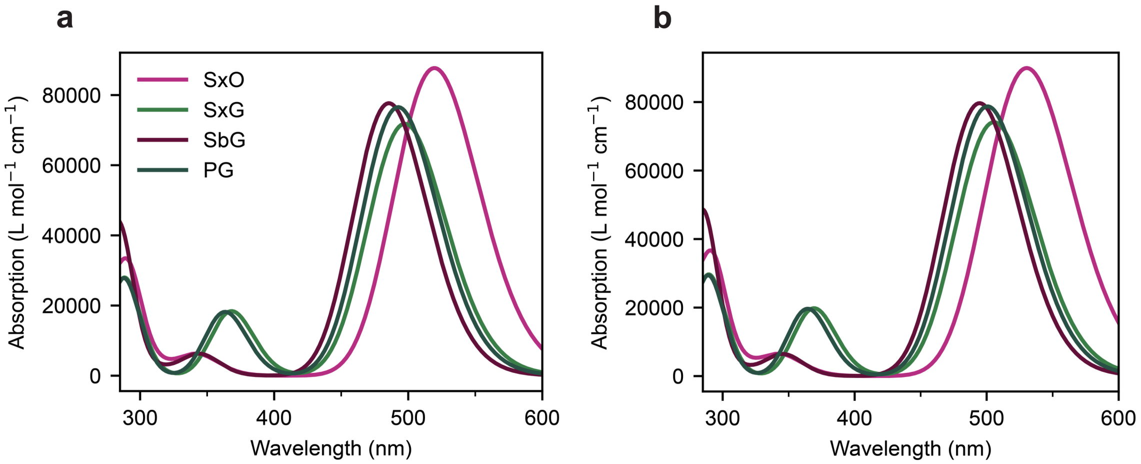


**Supplementary Figure S21. Simulated absorption spectra for SxO, SxG, SYBR Gold (SbG) and PicoGreen (PG).** The first ten excited states for each molecule were computed using RI-CC2/aug-cc-pVDZ employing the COSMO solvation model with a) DMSO and b) water as solvent. Absorption spectra were obtained by convoluting the calculated transitions with Gaussian functions with a full-width at half-maximum of 0.34 eV.
